# Elasticity control of entangled chromosomes: crosstalk between condensin complexes and nucleosomes

**DOI:** 10.1101/2022.11.09.515745

**Authors:** Tetsuya Yamamoto, Kazuhisa Kinoshita, Tatsuya Hirano

## Abstract

Condensin-mediated loop extrusion is now considered as the main driving force of mitotic chromosome assembly. Recent experiments have shown, however, that a class of mutant condensin complexes deficient in loop extrusion can assemble chromosome-like structures in *Xenopus* egg extracts, although these structures are somewhat different from those assembled by wild-type condensin complexes. In the absence of topoisomerase II (topo II), the mutant condensin complexes produce an unusual round-shaped structure termed a ‘bean’, which consists of a DNA-dense central core surrounded by a DNA-sparse halo. The mutant condensin complexes accumulate in the core whereas histones are more concentrated in the halo than in the core. We consider that this peculiar structure serves as a model system to study how DNA entanglements, nucleosomes, and condensin functionally crosstalk with each other. To gain insight into how the bean structure is formed, here we construct a theoretical model. Our theory predicts that the core is formed by attractive interactions between mutant condensin complexes whereas the halo is stabilized by the energy reduction through the selective accumulation of nucleosomes. The formation of the halo increases the elastic free energy due to the DNA entanglement in the core, but the latter free energy is compensated by condensin complexes that suppress the assembly of nucleosomes.

## INTRODUCTION

Upon entry into mitosis, chromosomal DNAs undergo a series of dramatic structural transitions from the relatively diffuse interphase chromatin structures to a discrete set of rod-shaped structures, each composed of duplicated sister chromatids. This process, collectively referred to as mitotic chromosome assembly, is essential for the faithful segregation of genetic information into daughter cells. Extensive studies over the past two decades or so have fully established that condensins, a class of SMC protein complexes, play a central role in this process (Hirano, 2016). Although it remains unknown exactly how condensins might work at a mechanistic level, the so-called loop extrusion theory has recently attracted much attention (Alipour and Marko, 2012, Naumova et al., 2013, Goloborodko et al., 2016a and 2016b). The theory originally predicted that the rod-shaped structure of mitotic chromosomes is assembled by the loop extrusion mechanism. The subsequent demonstration that budding yeast condensin has the activity to translocate along a DNA (Terakawa et al., 2017) and to extrude DNA loops (Ganji et al., 2018, Kong et al., 2020) in vitro has provided strong experimental support for this theory.

Is the loop extrusion mediated by condensins sufficient to assemble mitotic chromosomes? Early studies demonstrated that *Xenopus laevis* condensin I has the activity to introduce positive superhelical twist into double-stranded DNAs (Kimura and Hirano, 1997). Moreover, it is not fully understood how condensins extrude chromatinized DNA substrates or how they functionally cross-talk to other chromosomal components, such as topoisomerase II (topo II) and histones, during mitotic chromosome assembly (Shintomi et al., 2015; Shintomi et al., 2017, Choppakatia et al., 2021; Shintomi et al., 2021). Experiments using *Xenopus* egg extracts have shown that mutant condensin complexes deficient in loop extrusion are not completely defective in chromosome assembly (Kinoshita et al. 2015; Kinoshita et. al., 2022; see Fig. 1**a**). Notably, under the condition where DNA entanglements between different chromosomes persist in the absence of topo II, the addition of the mutant condensin complexes converts a swollen DNA network (Fig. 1**b**) into an unusually compact structure, called a ‘bean’, that is composed of a DNA-dense core and a DNA-sparse halo (Fig. 1**c**). The mutant condensin complexes accumulate in the core whereas histones are more concentrated in the halo than in the core. These studies raised the possibility that a loop extrusion-independent mechanism contributes to mitotic chromosome assembly and shaping. It has been proposed that condensin-condensin interactions through its HEAT repeat subunit could underlie these processes (Sakai et al. 2018; Kinoshita et al. 2022). Another example of a loop extrusion-independent mechanism of DNA compaction by SMC protein complexes was suggested through experiments using a mixture of bare DNAs and yeast cohesin complexes (Ryu et al. 2021). In this study, DNA compaction was explained by DNA-cohesin-DNA bridging (Brackley et al. 2013, Brackley et al. 2017).

**Figure 1.**
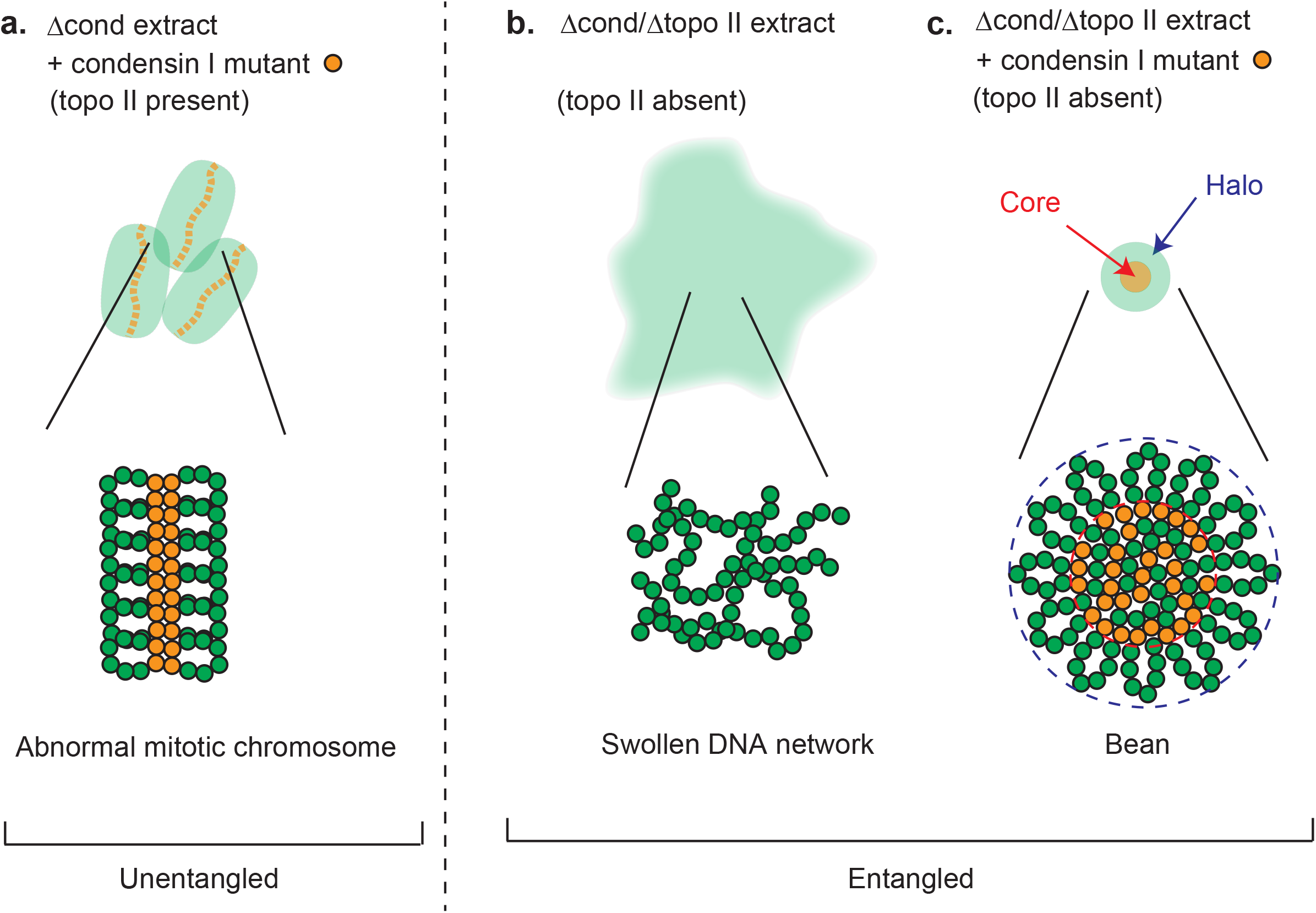
Polymorphism of chromatin structures observed in *Xenopus* egg extracts. Shown above are chromatin structures observed under different conditions in *Xenopus* egg extracts. Shown below are more detailed structures with mutant condensin I complexes deficient in loop extrusion (orange circles) and DNA units (green circles). **a**. Abnormal chromosome-like structures observed in a condensin-depleted extract (Δcond extract) that is supplemented with a condensin I mutant. **b**. A ‘cloud’-like structure observed in a condensin/topo II-depleted extract (Δcond Δtopo II extract). **c**. A ‘bean’ structure observed in a condensin/topo II-depleted extract (Δcond Δtopo II extract) that is supplemented with the condensin I mutant. Under the conditions of **b** and **c**, entangled DNAs behave as a polymer network.

Entanglements restrict the fluctuation of polymer chains and are treated as effective crosslinks in polymer physics (Edwards, 1967, Rubinstein and Colby, 2003). Unlike chemical crosslinks, these effective crosslinks can slide along the chains and are therefore referred to as slip-links (Ball et al., 1981, Masubuchi, 2021) or slip-springs (Likhtman, 2005, Uneyama and Masubuchi, 2012). In the absence of topo II, entanglements between different chromosomes are not resolved in the experimental time scales (Rosa and Everaers, 2008) and can therefore be treated as an entangled polymer network (Fig. 1**c**). In general, when the polymer network is stretched, a subset of chains in the network is stretched and contributes to the elasticity of the network (elastically effective chains; the black broken lines in Fig. 2**a**) (Rubinstein and Colby, 2003). The other chains such as dangling loops, which do not respond to the stretching of the network, do not contribute to the elasticity of the network (elastically ineffective chains; the light blue broken lines in Fig. 2**a**). When nucleosomes are assembled, DNA subchains wrapped around histones become elastically ineffective (the cyan broken lines in Fig. 2**b**). We also count the DNA loops in the halo of a bean elastically ineffective because these loops are treated separately from the network in the core (Fig. 2**c**). Polymer physics predicts that the stiffness of an elastically effective chain increases as the number of DNA units in the chain decreases (Rubinstein and Colby, 2003). The formation of DNA loops in the halo stiffens the entangled DNA network in the core by reeling DNAs out and decreasing the number of DNA units in the elastically effective part of DNAs. Recently, a similar concept was used to theoretically predict the contribution of the loop extrusion to the elasticity of entangled bare DNAs (Yamamoto and Schiessel, 2022). The electrostatic aspect of the nucleosome assembly, such as the charge inversion (Park et al., 1999, Nguyen and Shklovskii, 2001), is a classical subject of soft matter and biological physics. However, the regulation of DNA elasticity by functional crosstalk between nucleosomes and condensin complexes is only rarely studied, especially in the context of large-scale chromosome assembly.

**Figure 2.**
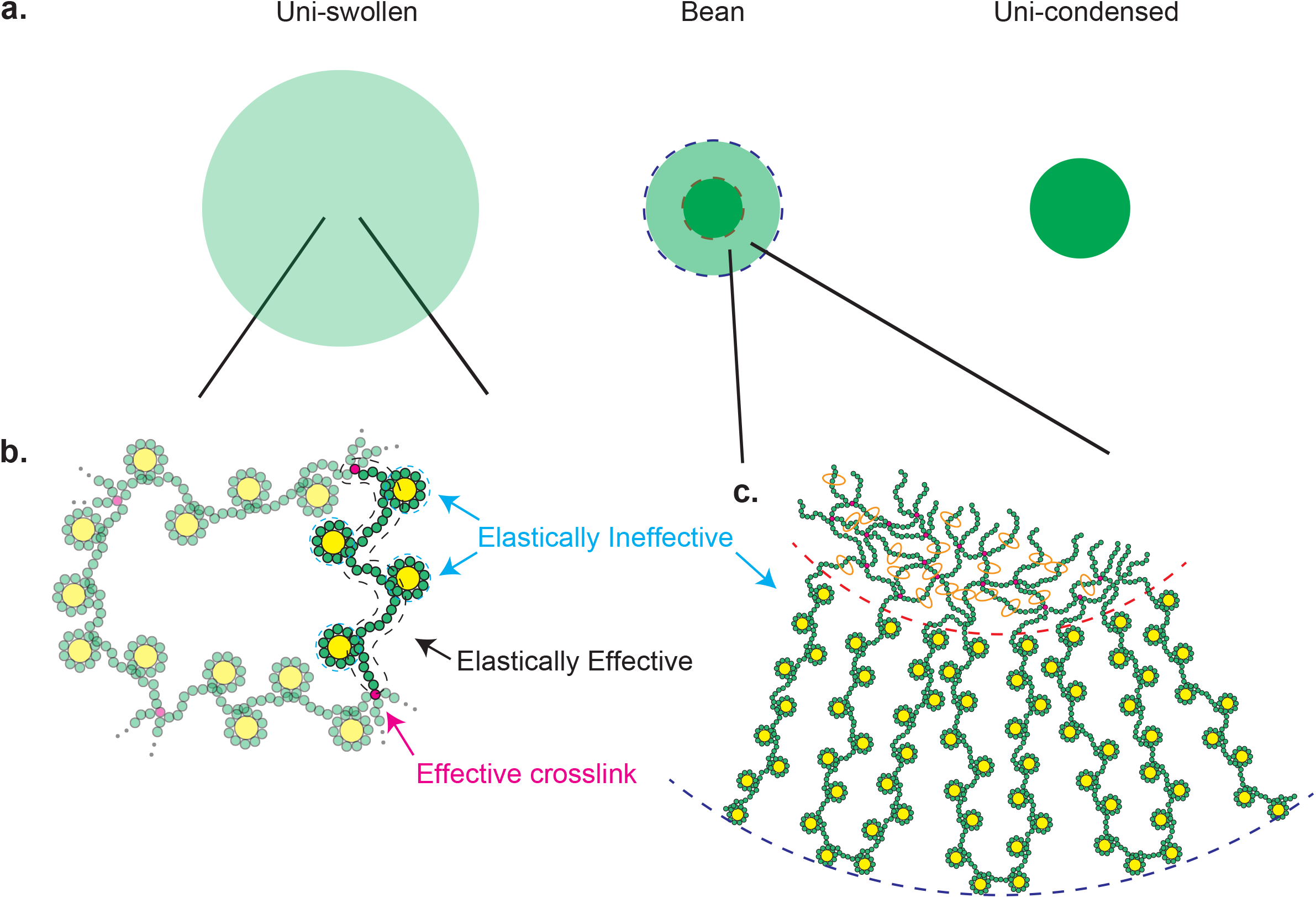
Elastically effective and ineffective chains in an entangled DNA network. **a**. Three possible structures formed by entangled DNAs: the all-swollen state (left), the bean structure (middle), and the un-condensed state (right). **b**. DNA chains connected to the network at two points via effective crosslinks due to DNA entanglements (magenta) respond to the deformation of the network and thus are elastically effective (indicated by the black broken lines). DNA units wrapped around histones (yellow circles) do not respond to such network deformation and thus are elastically ineffective (indicated by the cyan broken line). **c**. The DNA loops in the halo do not contribute to the elastic free energy due to the entanglement, thereby being elastically ineffective.

To provide insight into the postulated loop extrusion-independent mechanism of chromosome assembly, we here construct a mean field theory of entangled DNAs under the condition where mutant condensin complexes deficient in loop extrusion form the bean structure. This theory predicts that the attractive interaction between mutant condensin complexes drive the formation of the DNA-dense and nucleosome-poor central core of a bean whereas the DNA-sparse halo is stabilized by the free energy reduction associated with the assembly of nucleosomes. The formation of the halo in turn increases the free energy contribution of the DNA entanglement in the core, the free energy contribution of the excluded volume interaction between nucleosomal DNAs in the halo, and the elastic free energy by forming DNA loops in the halo. The free energy increase due to the entanglement is compensated by mutant condensin complexes that maintain the flexibility of the elastically effective chains in the core by suppressing the assembly of nucleosomes. Critical experimental tests on these predictions will provide insight into the biophysical properties of nucleosomal DNAs and condensin complexes involved in the assembly of mitotic chromosomes.

## MATERIALS AND METHODS

### Mechanical properties of bare and nucleosomal DNA

In an aqueous solution of physiological concentration, the electric charges of DNAs are neutralized by the Manning condensation and the Debye-Huckel screening due to salt ions (Marko and Siggia, 1995, Bracha et al., 2013). The unit length *b*_0_ of a bare DNA is 100 nm (≈ 300 bps) (Marko and Siggia, 1995). In a recent experiment on yeast, the unit length *b* of nucleosomal DNAs was estimated as 50 nm and there are 2 nucleosomes per 10 nm (Socol et al., 2019). A stretch of DNA is wound around an octamer of histone proteins by ≈146 bps in each nucleosome and nucleosomes are spaced by linker DNA regions of ≈ 19 bps: DNA of length 1.65 kbps is thus included in a Kuhn length of nucleosomal DNA. In this paper, we define a DNA unit as the DNA in a Kuhn length of bare DNA. *g*_n_ (≈ 5.5) is the number of DNA units in a Kuhn length of nucleosomal DNA. Chromosomes are much longer than either of the two Kuhn lengths, *b*_0_ and *b*, and are treated as electrically neutral flexible polymers.

### Model of the bean structure

In this theory, we theoretically predict the structure of entangled DNAs in an aqueous solution, including histone and the mutant condensin complexes deficient in loop extrusion (Fig. 3). We mainly treat the uniform polymer model in which these DNAs are assumed to be homogeneous polymers in the chromosome length scale. We treat the multi-block copolymer model in which the DNAs are assumed to be copolymers composed of two types of blocks with different free energy for nucleosome assembly only approximately and briefly. The bean structure is composed of the core and the halo. We model the core as a network of entangled DNAs and the halo as a brush of DNA loops at the surface of the core. A fraction of DNA units are distributed to the core, while the other units are distributed to the halo. The second law of thermodynamics states that the most stable state is given by the minimum of the free energy. In this theory, we seek the mechanism of the assembly of the bean structure by analyzing the minimum of the free energy.

**Figure 3.**
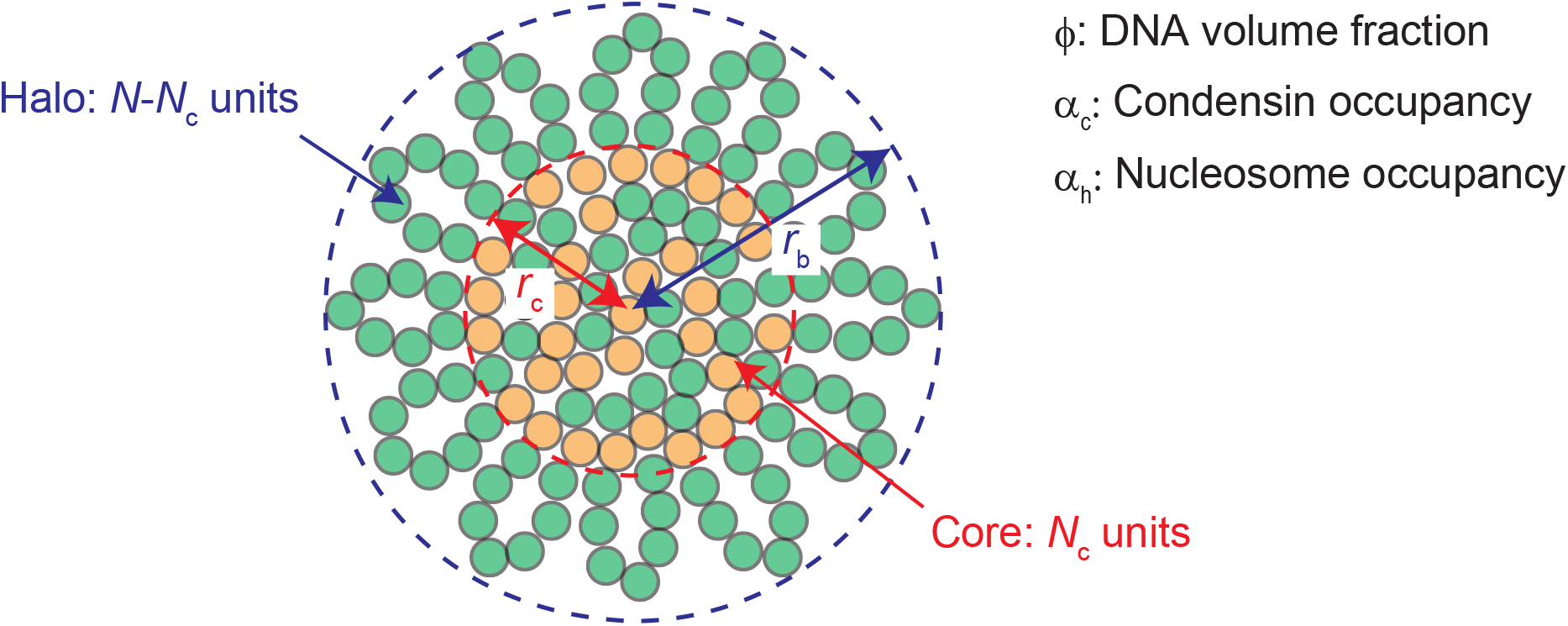
Model of the bean structure. The bean is composed of the core and the halo. The core and the halo are modeled as an entangled DNA network and a brush of DNA loops. Each DNA segment can form nucleosomes, can be occupied by mutant condensinsor can be bare. A fraction *N*_c_/*N* of DNA units is distributed in the core and the other DNA units are distributed in the halo. The bean is characterized by the DNA volume fraction *ϕ*, the occupancies, *α*_*c*_ and *α*_*h*_, of DNA by the mutant condensin complexes and nucleosomes.

The free energy *F* of the system is represented as the sum of the free energy *F*_c_ of the core and the free energy *F*_h_ of the halo,

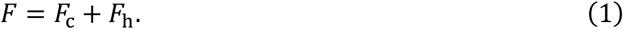

For simplicity, we use the mean field theory, an extension of the Flory-Rehner theory (Flory and Rehner, 1943) for the core and an extension of the Alexander theory (Alexander, 1977, de Gennes, 1980) for the halo, to derive the free energy of the core and the halo, and we limit our discussion to a spherical bean.

The free energy of the core has the form

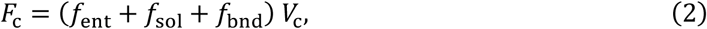

where *V*_c_ is the volume of the core. *f*_nt_ is the elastic free energy density due to the entanglement of DNAs in the core. *f*_sol_ is the solution free energy density, including the free energy contributions of the mixing entropy and the interactions between the mutant condensin complexes. *f*_bnd_ is the binding free energy density, including the contributions of the assembly of nucleosomes and the loading of the mutant condensin complexes. The structure of the core is quantified by the volume fraction *ϕ* of DNAs, the occupancies of DNA units by nucleosomes *α*_h_ and the mutant condensin complexes *α*_c_.

The free energy of the halo has the form

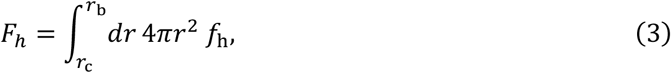

where *r* is the radial coordinate to represent positions in a bean structure by the distance from the center of the spherical bean. *r*_c_ is the radius of the core and *r*_b_ is the radius of the bean. The free energy density of the halo has three contributions

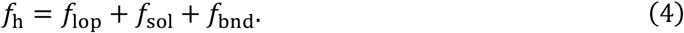

*f*_lop_ is the elastic free energy density to form DNA loops and is independent of the entanglement. *f*_sol_ is the solution free energy density and *f*_bnd_ is the binding free energy density. The free energy contributions, *f*_sol_ and *f*_bnd_, are the same as the second and third terms in the integrand of eq. (2). These free energy densities are functions of the local DNA volume fraction *ϕ*, the local occupancies of nucleosomes *α*_h_ and the mutant condensin complexes *α*_c_, which are functions of the coordinate *r*.

The free energy of the entangled DNAs does not depend on the length of each of the DNAs (Rubinstein and Colby, 2003). We thus specify the system by the number *N*_c_ of DNA units in the core and the number *N* − *N*_c_ of DNA units in the halo (*N* is the total number of DNA units in the system). The volume *V*_c_ and radius *r*_c_ of the core has a relationship with the DNA volume fraction *ϕ* in the core and the number *N*_c_ of DNA units in the core,

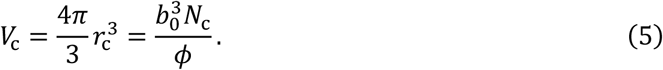

The radius *r*_b_ of the bean also has a relationship with the DNA volume fraction *ϕ* in the halo and the number *N* − *N*_c_ of DNA units in the halo

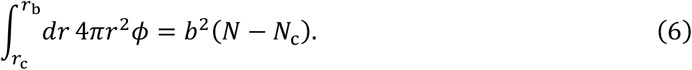

### Elastic free energy due to the entanglement

In polymer physics, the entanglement is treated as effective crosslinks. These effective crosslinks are distributed uniformly in the core, while there are no entanglements in the halo. We use the simplest model of polymer entanglement, the affine tube model that predicts that the effective crosslinks due to the entanglement are fixed at the space (more precisely, this theory takes into account the Gaussian fluctuation around the fixed point, but it does not change the physics) (Edwards, 1967, Rubinstein and Colby, 2003). In contrast to chemical crosslinks, chains can slide through the effective crosslinks. If we assume the DNAs are nucleosomal in the reference state (when these DNAs are entangled and thus are relaxed), the end-to-end distance of a DNA chain is

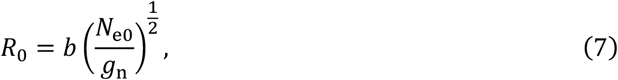

where *N*_e0_ is the number of DNA units in a subchain between two effective crosslinks in the relaxed state (count by bare DNA units). When the DNA network is swollen or deswollen so that the dimension of the network changes by *λ*_s_ times (*λ*_s_ is called the swelling ratio), the positions of effective crosslinks displace with the same ratio (the affine deformation), see Fig. 4. The end-to-end distance of the DNA chain thus changes to *λ*_*s*_*R*_0_.

**Figure 4.**
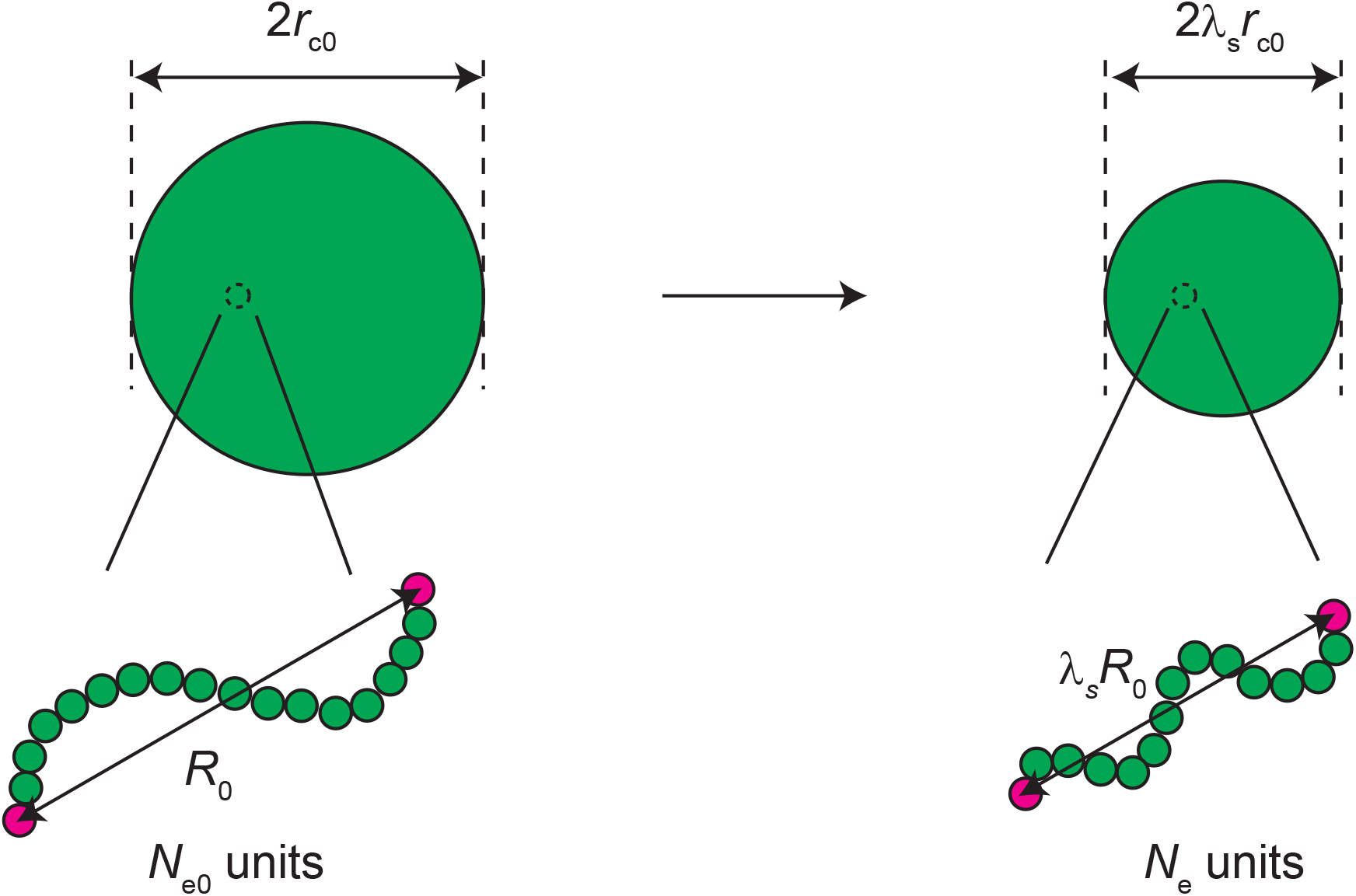
Affine deformation. In the relaxed (reference) state, the radius of the core is *r*_c0_ and the average distance between the two effective crosslinks that connect a DNA chain composed of *N*_e0_ is *R*_0_. If the core is swollen or deswollen to the radius *r*_c_ (= *λ*_s_*r*_c0_), the average distance between the effective crosslinks changes to *λ*_s_*R*_0_ (affine deformation). If the halo is assembled, the number of DNA units in the chain between the effective crosslinks changes to *N*_e_.

In the reference state, all DNA units are distributed to the core. In the bean structure, some DNA units are distributed to the halo and thus the number of DNA units in the core decreases from that of the reference state. DNA units are distributed to make the tensions of chains between effective crosslinks equal. Because the number of effective crosslinks is constant, the number *N*_e_ of DNA units between two effective crosslinks in the bean structure has the form

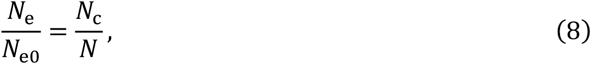

where *N*_c_ is the total number of DNA units in the core and *N* is the total number of DNA units in the system. If the DNA chain is bare, the square *δR*^2^(*N*_e_) of the amplitude of the fluctuations of the chain is 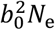, whereas, if the DNA chain is fully packed with nucleosomes, *δR*^2^(*N*_e_) is *b*^2^*N*_e_/*g*_n_. We use the linear interpolation

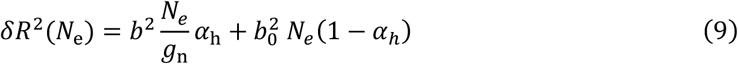

to treat cases in which the occupancy of DNA units by nucleosomes is *a*_h_. We take into account the fact that DNA units wound around histone proteins and those at the halo are elastically ineffective via eq. (9). The linear approximation, eq. (9), corresponds to the serial connection of a spring of bare DNA and a spring of nucleosomal DNA. This approximation is exact if we only take into account the changes in the number of DNA units in the elastically effective part of the chain, *b* → *b*_0_ and *g*_n_ → 165 bps/19 pbs ≈ 8.7. In the treatment of eq. (9), we also take into account the difference in the Kuhn length between nucleosomal DNAs and bare DNAs up to the linear term of *α*_h_.

The volume fraction of DNA units is *ϕ*_0_ in the reference state. In the core of the bean, the volume fraction of DNA units changes to

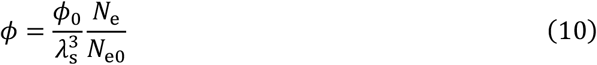

because the volume of the core changes by 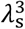 by the swelling or deswelling and the number of DNA units in the core decreases to *N*_e_ because of the assembly of the halo.

The elastic free energy density is the elastic energy of each chain 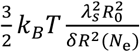, multiplied by the volume density *ϕ*/(*b*^3^*N*_e_) of DNA in the network (Rubinstein and Colby, 2003, Yamamoto et al., 2022**a**),

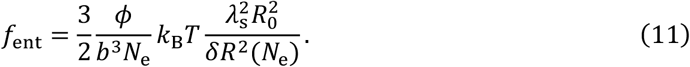

Eq. (11) is simplified to

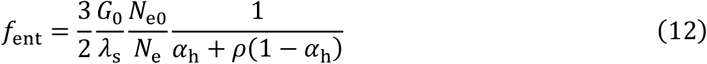

by using eq. (10) and the shear modulus of the network at the reference state

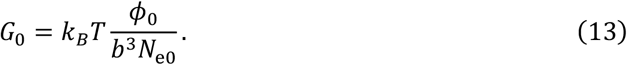

The factor *ρ* in eq. (12) is the ratio of the squared amplitudes of fluctuations between the bare and nucleosomal DNA with the same DNA length,

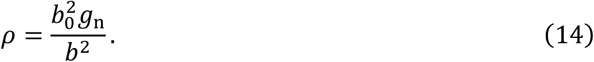

The factor *N*_e0_/*N*_e_ in eq. (12) represents the stifening of the entangled DNAs in the core by reeling DNA units from the core to the halo. The factor 1/(*a*_h_ + *ρ*(1 − *a*_h_)) in eq. (12) represents the stifening of the entangled DNAs in the core by assembling nucleosomes.

### Solution free energy

The solution free energy includes the contribution of the mixing entropy and the contribution of the condensin-condensin interactions. The conformational transitions of condensin complexes influence the interactions between these complexes (Kinoshita et al., 2015). The key feature of the conformational transitions, revealed by recent X-ray crystallography (Hassler et al., 2019) and cryo-EM (Lee et al., 2020), is that the SMC4 subunit of a condensin complex binds to the SMC2 subunit in the presence of ATP whereas it binds to CAP-D2 in its absence (a flip-flop mechanism). It has been proposed that a condensin-condensin interaction may be triggered by a contact between SMC4 in one complex with CAP-D2 in another complex in the absence of ATP (Kinoshita et al., 2022).

The kinetics of the fraction *θ* of the mutant condensin complexes, to which ATPs are not bound and thus interact with other condensin complexes, is represented by the time evolution equation

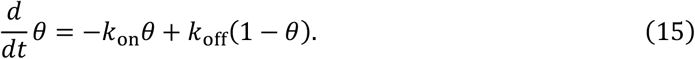

In the steady state, the fraction *θ* is derived as

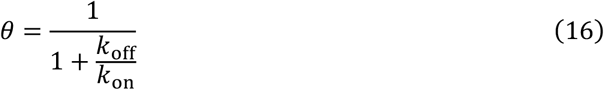

by using eq. (15). In our model, we assume that the condensin-condensin interaction is multivalent, like other theories of mitotic chromosome assembly (Sakai et al. 2018) and transition (Forte et al. 2022). The free energy density due to the interaction between condensin complexes has the form

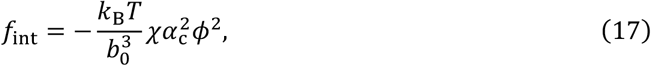

where *χ* is the magnitude of the attractive interaction between the mutant condensin complexes. The interaction parameter *χ* has the form

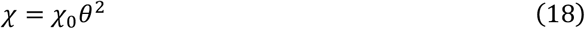

because ATPs are not bound to both of the interacting condensin complexes. With eq. (17), the solution free energy density has the form

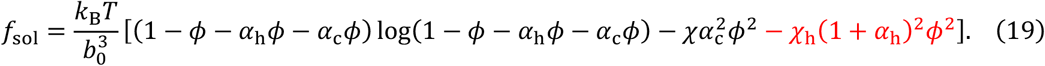

This free energy, eq. (19), is composed of the mixing free energy (the first term), the free energy of the condensin-condensin interaction (the second term), and the free energy contribution of the DNA-DNA interaction (the third term). The mixing free energy quantifies the random thermal motion that mixes DNA units and solvent and acts as the effective excluded volume interaction between DNA units (Rubinstein and Colby, 2003). The swelling of polymer networks is driven by this effective excluded volume interaction. The factor *α*_h_*ϕ* represents the increase of the excluded volume due to the assembly of nucleosomes and the factor *α*_c_*ϕ* is the increase of excluded volume due to the loading of condensin complexes. For simplicity, we assume that for both cases, the increase of the excluded volume is the same as the volume of DNA units. The free energy contribution of the excluded volume interaction is included in the first term of eq. (19). *χ*_h_ is the interaction parameter that accounts for the DNA-DNA attractive interaction. We study the athermal solvent condition in which the magnitude of the two-body DNA-DNA excluded volume interaction is maximum (*χ*_h_ = 0) and the θ-solvent condition in which the excluded volume interaction is negligible (*χ*_h_ = 1/2) with the mixing free energy, see the first term of eq. (19).

### Binding free energy

The binding free energy has the form

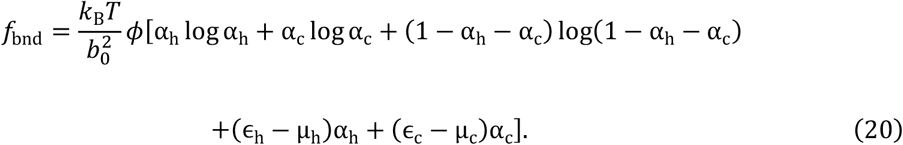

The first, second, and third terms of eq. (20) are the free energy contributions of the entropy with respect to the assembly of nucleosomes and the loading of the mutant condensin complexes. With this treatment, we assume that nucleosomes and the mutant condensin complexes cannot occupy the same DNA unit simultaneously, *α*_h_ + *α*_c_ ≤ 1. The chemical potentials, 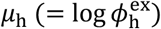 and 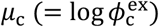 are the logarithms of the volume fractions, 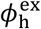 and 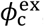, of histones and the mutant condensin complexes in the solution at the exterior to the bean. The fourth term is the energy gain *k*_B_*Tϵ*_h_ due to the assembly of a nucleosome, relative to the chemical potential *k*_B_*Tµ*_h_ of histone proteins. The fifth term results from the stabilizing energy *k*_B_*Tϵ*_c_ due to the loading of the mutant condensin complexes, relative to the chemical potential *k*_B_*Tµ*_c_ of these mutants. Our theory only analyze the structure of entangled DNAs at the steady state and thus does not explicitly take into account the diffusion of mutant condensins along DNAs.

### Elastic free energy of DNA loops

The DNAs in the halo are treated as a brush of DNA loops. The simplest treatment of a polymer brush is the Alexander model which assumes that the concentration of polymer units is uniform in the brush for the limit of vanishing curvature (Alexander, 1977, de Gennes, 1980). The DNA loops can be divided into ‘blobs’ composed of *g* DNA units, which occupy the volume *ξ*^3^, where *ξ* is the average distance between DNA chains at the same distance *r* from the center (de Gennes, 1980). The effect of the curvature on the elastic free energy is taken into account in the spirit of the Daoud-Cotton scaling theory (Daoud and Cotton, 1982). Because the spherical surface at a distance *r* is occupied by the blobs, the average distance *ξ* has a relationship

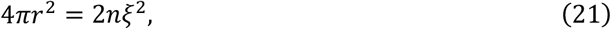

where *n* is the number of DNA loops and 2 in front of *n* represents the fact that each DNA loop contributes two DNA chains to the area. For simplicity, we assume that *n* is a constant. This treatment may be easily accepted in the multi-block copolymer model, where B blocks are preferentially localized in the halo and A blocks are preferentially localized in the core. In this model, *n* corresponds to the total number of B blocks in the entangled DNAs. The uniform chain model is the limit where the number of DNA units in each B block is small.

The elastic energy of the DNA loops in the halo thus has the form

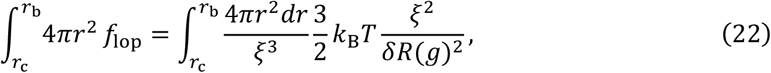

because the elastic free energy of each blob is 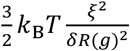 and 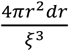 is the number of blobs at the distance *r*, see also eq. (9). The volume fraction of DNA units in the halo has the form

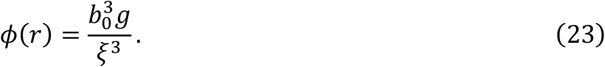

By using eqs. (21) and (23), eq. (22) is rewritten as a functional of *ϕ*(*r*) and *r*. This leads to the elastic free energy density of the loops in the form

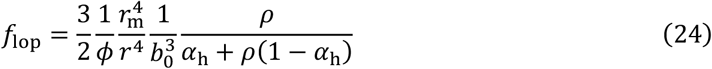

with

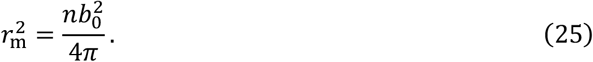

In the scaling theory of polymer brushes, the relationship between *ξ* and *g* is derived by the Flory theory, which corresponds to using eq. (44) (or just assumed the relationship *a priori*), and the free energy used for other purposes is estimated by *k*_B_*T* per blob (de Gennes, 1980). For simplicity, we here use eq. (44) for all purposes, although the scaling theory is quantitatively a better approximation.

### Multi-block copolymer model

In the multi-block copolymer model, we treat the entangled DNAs as multi-block AB copolymers, where the free energy *ϵ*_hA_ of assembling nucleosomes in the A blocks is larger than the free energy *ϵ*_hB_ of nucleosome assembly in the B blocks. The B blocks are thus preferentially localized at the halo by assembling nucleosomes, whereas the A blocks are preferentially localized to the core by loading mutant condensins. A fraction *N*_A_/*N* of DNA units is in the A blocks and the other fraction 1 − *N*_A_/*N* is in the B blocks. The free energy densities, *f*_c_(*α*_h_, *α*_c_, *ϕ*; *ϵ*_h_) and *f*_h_(*α*_h_, *α*_c_, *ϕ, r*; *ϵ*_h_), in the core and the halo are both functionals of the DNA volume fraction *ϕ* and occupancies, *α*_h_ and *α*_c_. We use the approximate free energy

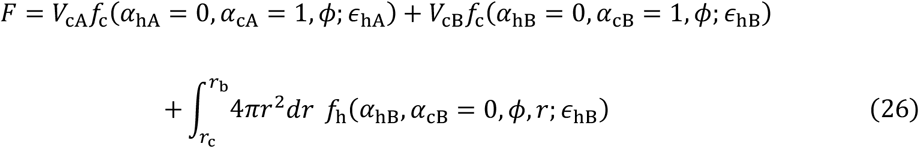

for *N*_c_ > *N*_A_ and

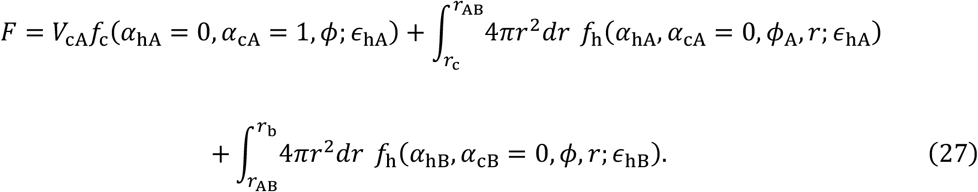

for *N*_c_ < *N*_A_. In eqs. (26) and (27), the nucleosome occupancy at the DNA units in the core and the condensin occupancy in the halo are neglected. This approximation enables us to avoid the complication arising from the fact that the deformation in the core is anisotropic and inhomogeneous for *N*_c_ > *N*_A_ because the extension ratio in the tangential direction to the interface between the A-rich and B-rich regions is continuous (Sekimoto, 1993, Tomari and Doi, 1995). For *N*_c_ < *N*_A_, the osmotic pressure is continuous at the interface between A-rich and B-rich regions, *r* = *r*_AB_.

### Parameter estimate

Our theory predicts the DNA volume fraction *ϕ*, the occupancies, *α*_h_ and *α*_c_, of nucleosomes and the mutant condensin complexes, and the fraction *N*_c_/*N* of DNA units in the core as functions of the rescaled shear modulus 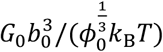, the free energy *ϵ*_h_ − *µ*_h_ gained by assembling nucleosomes, the free energy *ϵ*_c_ − *µ*_*c*_ gained by loading the mutant condensin complexes, the interaction parameter *χ*, the ratio *ρ* of the size of bare DNAs to nucleosomal DNAs (see eq. (14)), and a parameter *ϕ*_m_ defined by

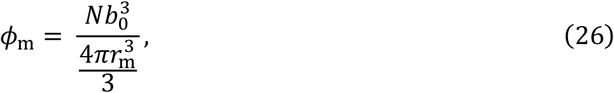

see eq. (25) for the definition of *r*_m_. The values of the parameters used for the numerical calculation are listed in Table 1.

**Table 1.**
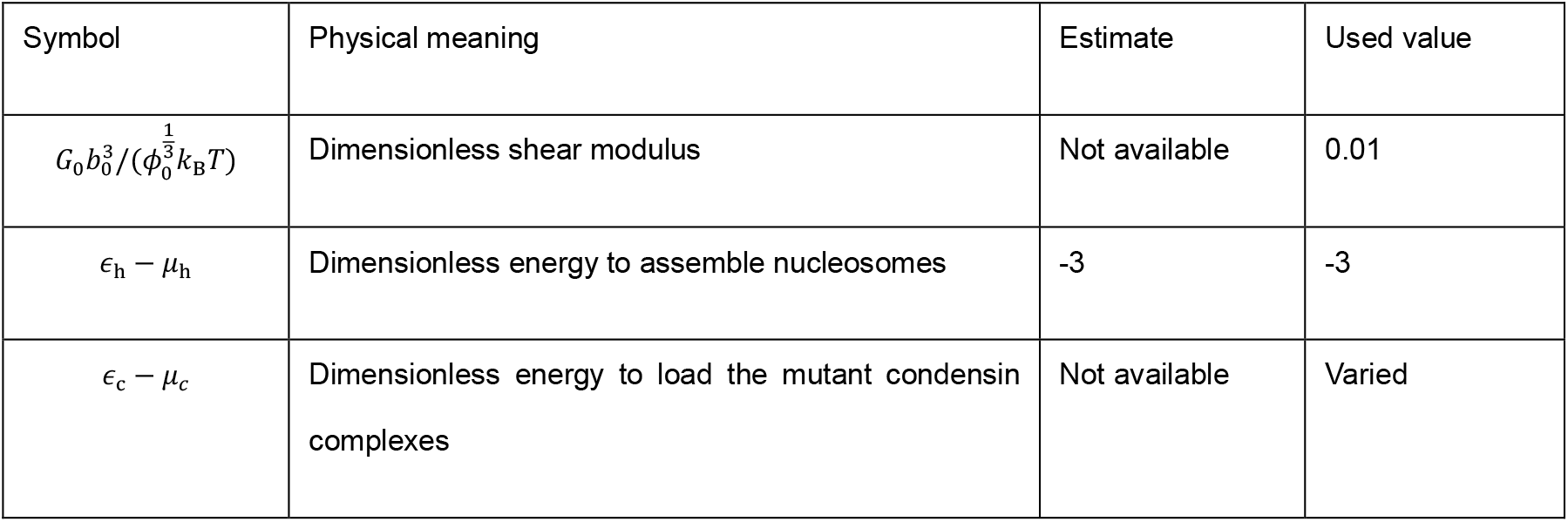
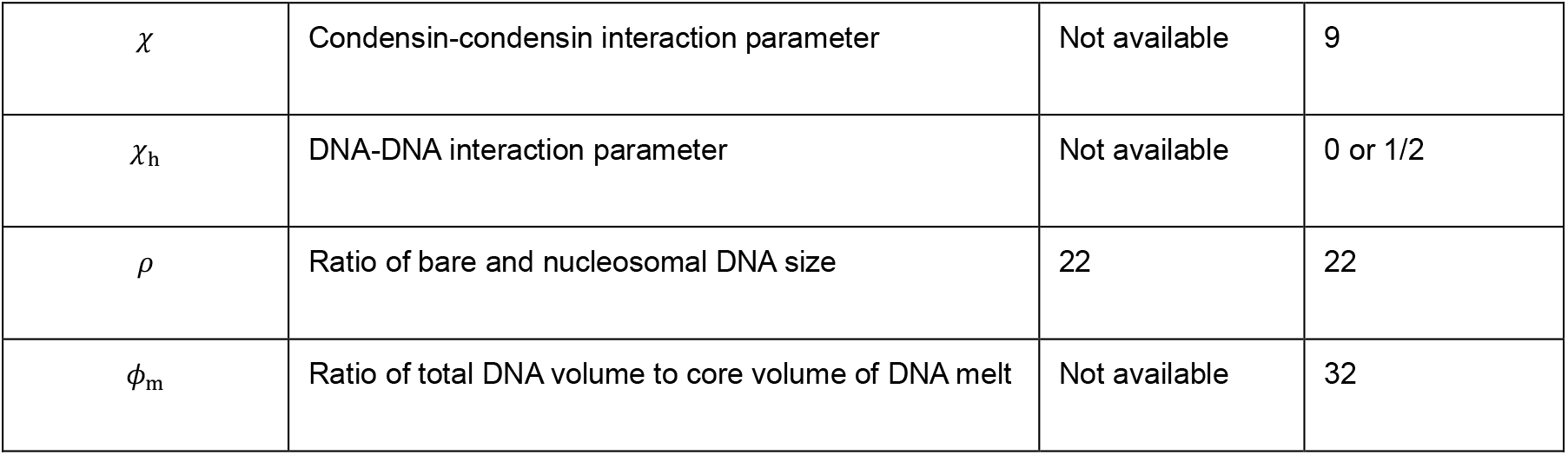
Values of parameters used for numerical calculations

The chemical potential of histone proteins is estimated as -12 by using the histone concentration 7 µM in the *Xenopus* egg extract and a rough estimate of the volume of histone proteins 1 nm^3^. This estimate assumes that histone octamers are assembled prior to the assembly of nucleosomes. The energy gained by assembling nucleosomes is estimated as -15 (Blossey and Schiessel, 2011). In principle, the rescaled shear modulus 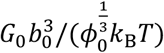 can be extracted by measuring the force-extension relationship of the entangled chromosome though it has not been characterized. We here choose the value of the parameter *ϕ*_m_ so that the radius *r*_b_ of the bean can be approximately twice lager than the radius *r*_c_ of the core as observed experimentally (Kinoshita et al. 2022).

## RESULTS

### The bean structure is stabilized by the free energy gained by assembling nucleosomes

For simplicity, we first use the uniform chain model to study the structure of entangled DNAs in an athermal solvent (*χ*_h_ = 0). The DNA volume fraction *ϕ*, the occupancies, *α*_h_ and *α*_c_, of DNA units by nucleosomes and the mutant condensin complexes, and the fraction *N*_c_/*N* of DNA units in the core of the stable structure are derived by using the conditions that the first derivatives of the free energy *F* with respect to these parameters are zero. These conditions correspond to the fact that the chemical potentials of DNA units, histone, and the mutant condensin complexes are uniform throughout the system and that the osmotic pressure is continuous at *r* = *r*_c_ and *r* = *r*_b_, where the halo interfaces with the core and the external solution, respectively, see also sec. S1 in the Supplementary Materials. By using these equations, we found a stable solution with which mutant condensins are localized at the core and nucleosomes are localized at the halo, in a window of the mutant condensin concentration, see Fig. 5. This solution will be called the bean state because the features of this solution are consistent with the experimentally observed bean structure. In the bean state, the free energy of nucleosomal assembly (the fourth term of eq. (20)) is dominant at the halo, while the free energy due to the condensin-condensin interaction (the second term of eq. (19)) and the free energy energy of mutant condensin loading (the fifth term of eq. (20)) are dominant at the core, see also eqs. (S38) and (S53) in the Supplemental Materials. The core is therefore stabilized by the attractive condensin-condensin interaction. This mechanism of core assembly is essentially the same as the bridging-induced phase separation (Brackley et al. 2013, Brackley et al. 2017, Forte et al. 2022). In contrast, the halo is stabilized by the free energy gain associated with nucleosome assembly.

**Figure 5.**
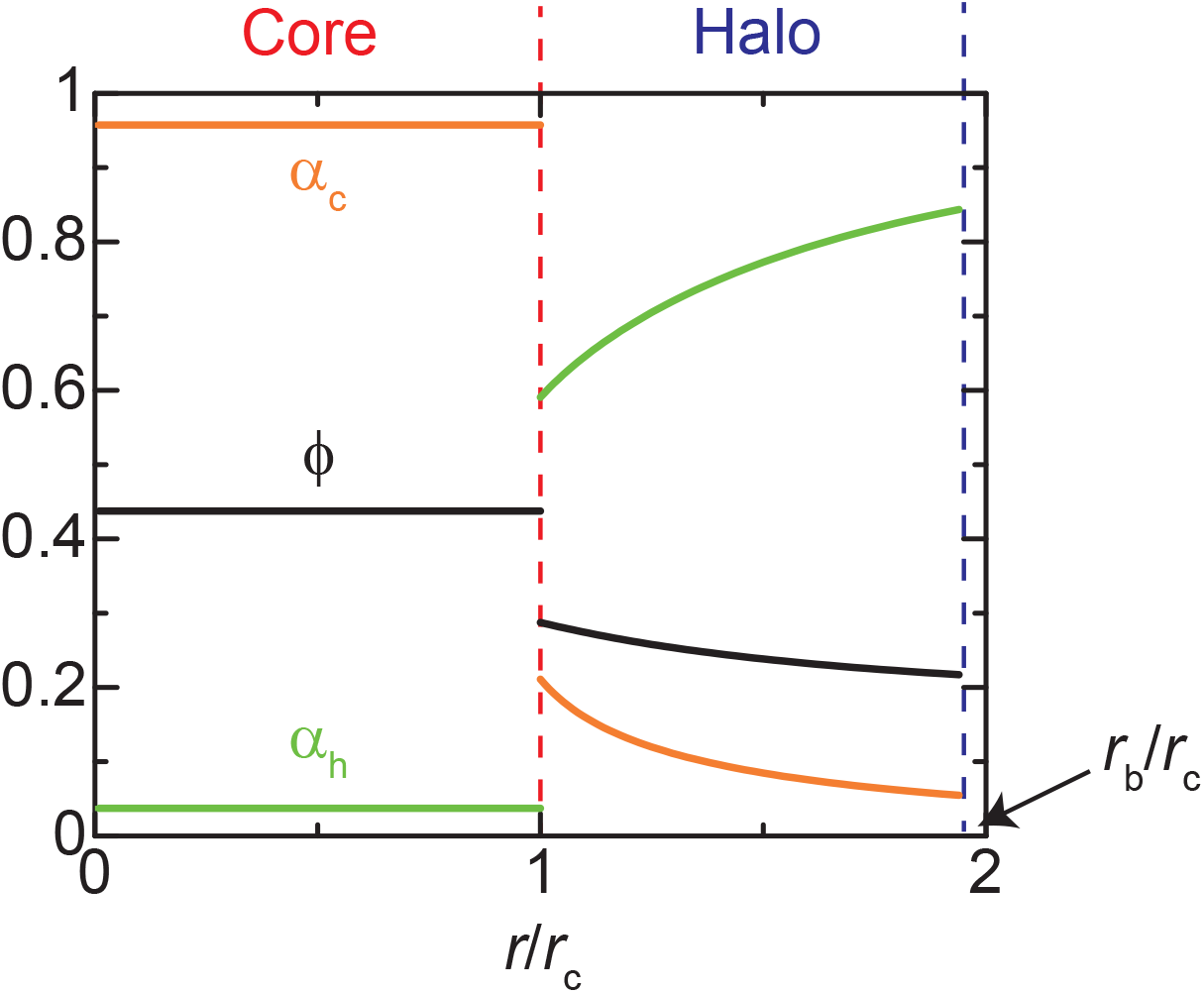
Profiles of DNA volume fraction and occupancies of DNA by the mutant condensin complexes and nucleosomes. The DNA volume fraction (black) and the occupancies, *α*_c_ and *α*_h_, of DNA by the mutant condensin complexes (orange) and nucleosomes (light green). The values of parameters used for the calculation are summarized in Table 1. We used *µ*_c_ − *ϵ*_c_ = -1.3 for the chemical potential of the mutant complexes.

### DNAs are reeled into the core as the concentration of the mutant condensin complex increases

Our theory predicts the radii, *r*_c_ and *r*_b_, of the core and the bean, see eq. (5) and (6). The radius *r*_c_ of the core increases, while the radius *r*_b_ of the bean decreases, with increasing the mutant condensin concentration, consistent with experiments (Kinoshita et al., 2022), see Fig. 6**a**. The condensed state of DNAs in the core becomes more stable as the concentration of the mutant condensin complexes increases. The fraction of DNA units distributed to the core increases with increasing the condensin concentration to gain the free energy due to the condensin-condensin interaction, see Fig. 6**b**. The sliding capability of the effective crosslinks enables the exchange of DNA between the core and the halo. For the condensin concentrations larger than a threshold value 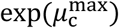, the bean structure is no longer stable and the entangled DNAs are uniformly condensed due to the condensin-condensin interaction (uni-condensed, Fig. 2**a**, right).

**Figure 6.**
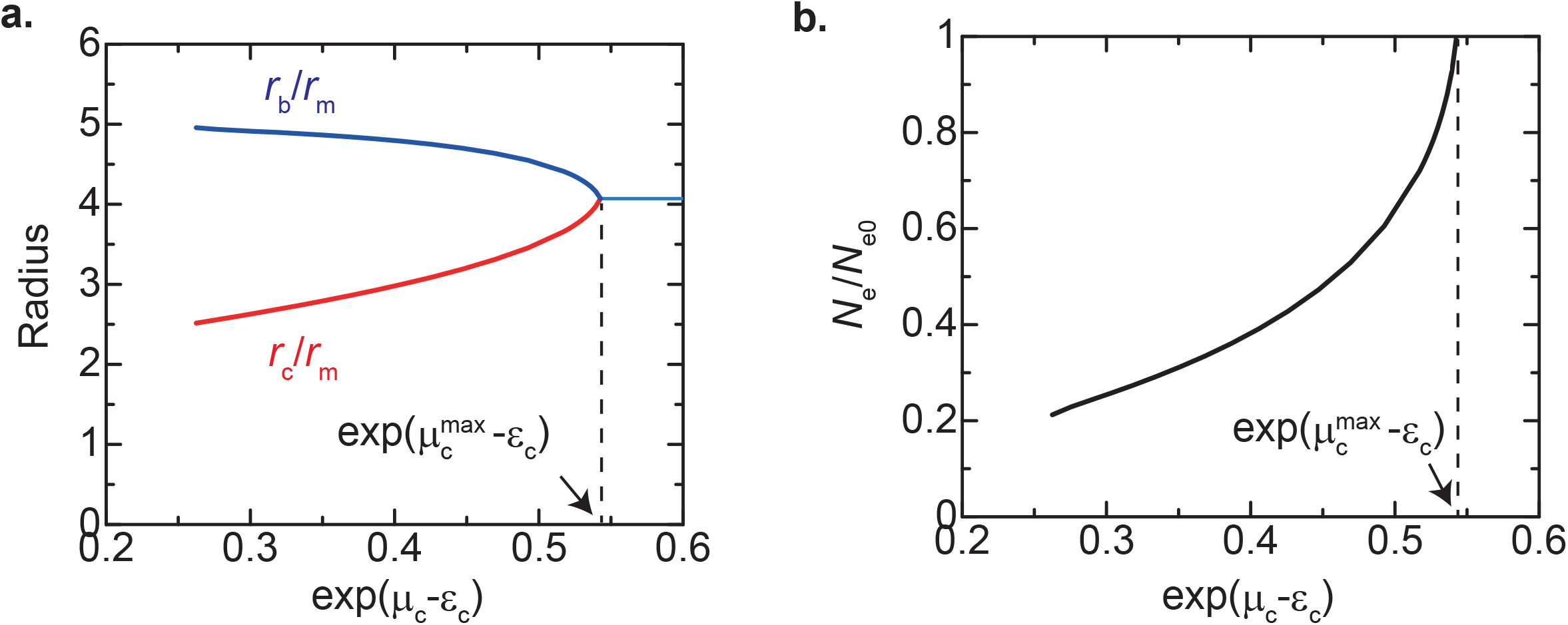
DNA is reeled into core as the concentration of the mutant condensin complexes increases. The radii of the core *r*_c_ (red) and the bean *r*_b_ (blue) (**a**) as well as the fraction of DNA units in the core (**b**) are shown as functions of the exponential exp(*µ*_c_ − *ϵ*_c_), which is proportional to the concentration of mutant condensins in the exterior solution. The parameters used for the calculation are summarized in Table 1.

### Uniform polymer model predicts that the bean structure is observed only if entangled DNAs are compacted by mutant condensins before nucleosomes are assembled

The fact that the halo is stabilized by the free energy gain associated with the assembly of nucleosomes implies that entangled DNAs are more stable if nucleosomes are assembled at the entire region of the DNAs (uni-swollen, Fig. 2**a**, left). Indeed, such a state satisfies the conditions of the free energy minimum (namely, the first derivatives of the free energy are zero). The free energy of the uni-swollen state is indeed smaller than the free energy of the bean state, implying that the bean structure is a local free energy minimum, but not the global free energy minimum, see Fig. 7. The bean structure is entrapped to the local free energy minimum by the free energy contributions of the condensin-condensin interaction and the loading of mutant condensins stabilizes the condensed core (see eq. (S38) in the Supplemental Materials). The entangled DNA network shows the bean structure if one decreases the concentration of the mutant condensin complexes from the uni-condensed state, but not if one increases the concentration of the mutant condensin complexes from the uni-swollen state. Our theory therefore predicts that the experimentally observed bean structure was kinetically trapped or there is an additional mechanism that further stabilizes the bean structure.

**Figure 7.**
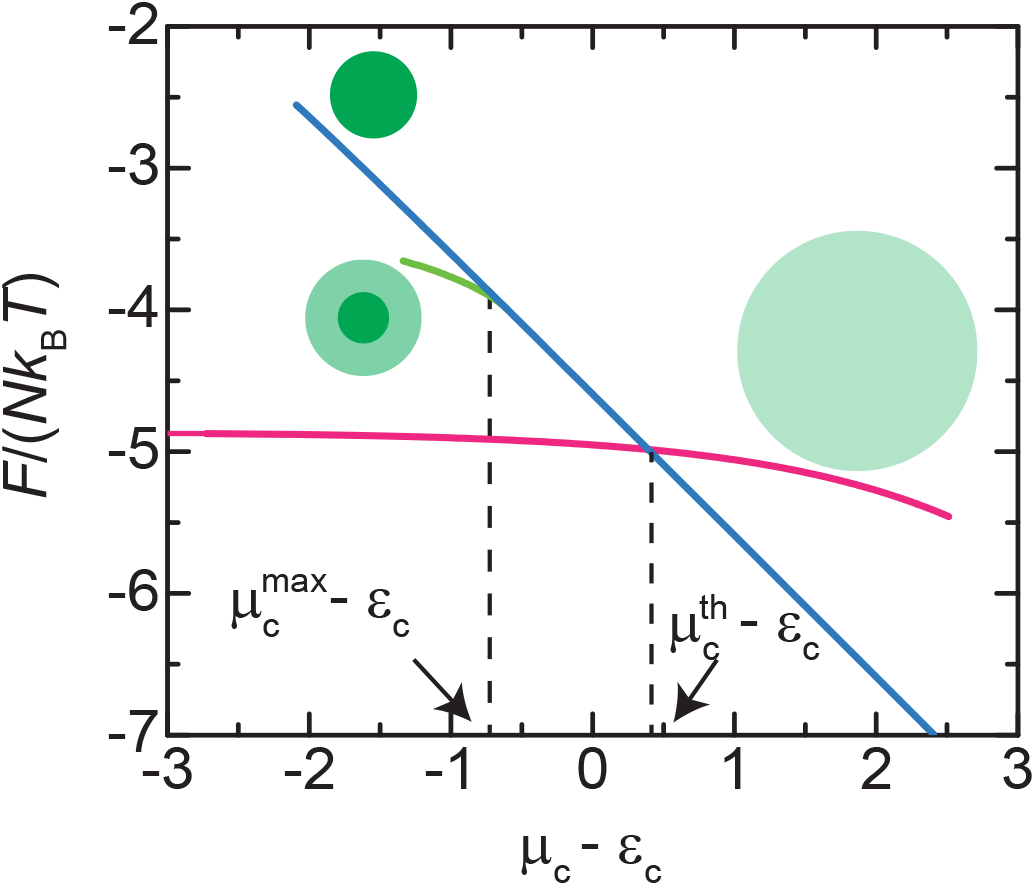
Free energy vs chemical potential of the mutant condensin complexes. The free energy of the system is shown as a function of the chemical potential of the mutant condensin complexes for the bean state (green), the uni-condensed state (cyan), and the uni-swollen state (magenta). The parameters used for the calculation are summarized in Table 1.

### Condensin complexes suppress nucleosome assembly and maintain the flexibility of elastically effective chains in the core

Among parameters used in our theory, the interaction parameter *χ* and the rescaled shear modulus 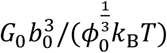 have not been experimentally determined. One may think that there may be a parameter space, in which the bean structure becomes the global free energy minimum. To be the global free energy minimum, the bean structure has to be stabilized up to the concentration of the mutant condensin complexes at the transition between uni-swollen and uni-condensed states, see Fig. 7. The maximum chemical potential 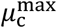 of the mutant condensin complexes, with which the bean structure is stable, and the chemical potential 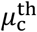 at the swollen-condensed transition are therefore the key quantity to discuss the stability of the bean structure (the following discussion is better presented by using the chemical potential, 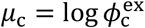, than the condensin concentration 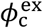 of the external solution). To understand the dependence of the stability of the bean structure on parameters explicitly, we derived an approximate analytical solution. With this approximation, we use *α*_c_ ≈ 1, *α*_h_ ≈ 0, and *ϕ* ≈ 1/2 for the core and *α*_c_ ≈ 0 for the halo. We expand the free energy in a power series of *ϕ* and *α*_h_ at *r* = *r*_c_ and neglect higher order terms, see sec. S2 for the details of the calculations.

The chemical potential 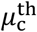 of the mutant condensin complexes at the swollen-condensed transition has an approximate form

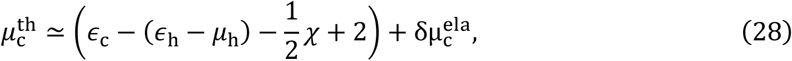

where 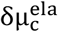 is the contribution of the entanglement free energy in the core and is a function of 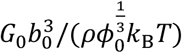, see eq. (S49) in the Supplemental Material. The maximum chemical potential of the mutant condensin complexes of the bean structure has an approximate form

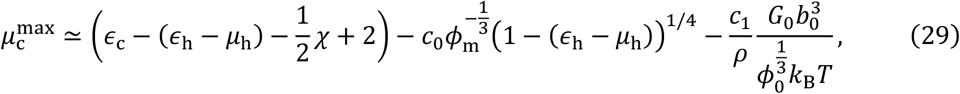

where *c*_0_ (≃2.6) and *c*_1_ (≃0.8) are numerical constants of order unity, see eq. (S65) in the Supplementary Materials. Eqs. (28) and (29) were derived in the asymptotic limit of large negative values of *ϵ*_h_ − *µ*_h_, although this approximation is somewhat crude for the parameters used for our numerical calculations, see sec. S2 in the Supplemental Materials. The first term of eq. (29) is the contribution of the interaction parameter *χ*, the free energy *ϵ*_h_ − *µ*_h_ gained by assembling nucleosomes, and the free energy *ϵ*_c_ gained by loading condensin complexes. The second term of eq. (29) is the contribution of the elastic free energy and the interaction free energy of the DNAs in the halo. The third term of eq. (29) is the contribution of the elastic free energy of entangled DNAs.

The two chemical potentials, 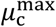 and 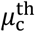, have the same dependence on *χ, ϵ*_h_ − *µ*_h_, *ϵ*_c_, implying that these parameters do not contribute to the stability of the bean structure, relative to the uni-swollen state, at least, in the leading order contributions, see the first terms of eqs. (28) and (29). The second and third terms of eq. (29) suppress the assembly of the halo, see their negative sign. The third term of eq. (29) is indeed not significant because of the large value of the ratio *ρ* ≃ 22 (while the rescaled shear modulus 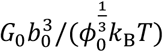 is smaller than 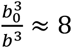). This implies that the elastic free energy of entangled DNAs does not increase significantly with the assembly of the halo because mutant condensins suppress the assembly of nucleosomes and maintain the number of DNA units in the elastically effective chains in the core, see eq. (11) and Table 1 (but it can still significantly destabilize the uni-swollen state via 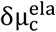, see Supplemental Material). The main mechanism that destabilizes the bean structure is the entropic elasticity of nucleosomal DNAs in the halo and the excluded volume interaction between these DNAs, see the second term of eq. (29).

At a first glance, one might think that the magnitude of the DNA-DNA excluded volume interaction is too large in the athermal solvent condition (*χ*_h_ = 0). However, even in the θ-solvent condition (*χ*_h_ = 1/2), where the two-body DNA-DNA excluded volume interaction is negligible, the bean structure is still a local free energy minimum, see Fig. S1. Note that the mean-field approximation is exact in the θ-solvent condition, implying that our results are not the artifacts of using the mean-field approximation.

### The bean structure may represent the global minimum if DNAs are multi-block copolymers

We have shown that the bean structure is not the most stable structure as long as we treat DNAs by using the uniform polymer model. Are there any other mechanisms that might help stabilize the bean structure? We consider that the bean structure may be analogous to the core-shell structure characteristic of paraspeckles. Paraspeckles are nuclear bodies scaffolded by NEAT1_2 architectural RNAs (arcRNAs), whicht form ribonucleoprotein (RNP) complexes with RNA-binding proteins (RBPs) (Clemson et al. 2009, Sasaki et al. 2009, Sunwoo et al. 2009). Recent studies have shown that NEAT1_2 RNP complexes act as ABC tri-block copolymers because different RBPs bind to the middle region and the terminal regions of NEAT1_2 arcRNAs; paraspeckles can be considered as micelles of such copolymers (Yamazaki et al. 2018 and 2021, Yamamoto et al. 2022**b**). The RBPs bound to each RNA region are determined by the local base sequence. The similarity between beans and paraspeckles motivated us to assume DNAs as multi-block AB copolymers, where the free energy *ϵ*_hA_ for nucleosome assembly in the A blocks is larger than the free energy *ϵ*_hB_ for nucleosome assembly in the B blocks (multi-block copolymer model), see Fig. 8**a** and also MATERIALS AND METHODS for the details of this model. In this model, it is not possible to discuss the limit of the instability of uni-condensed and uni-swollen phases, but it is sufficient to discuss the free energy in the window of multant condensin concentrations, where the bean structure is a local free energy minimum, see also Fig. S2.

**Figure 8.**
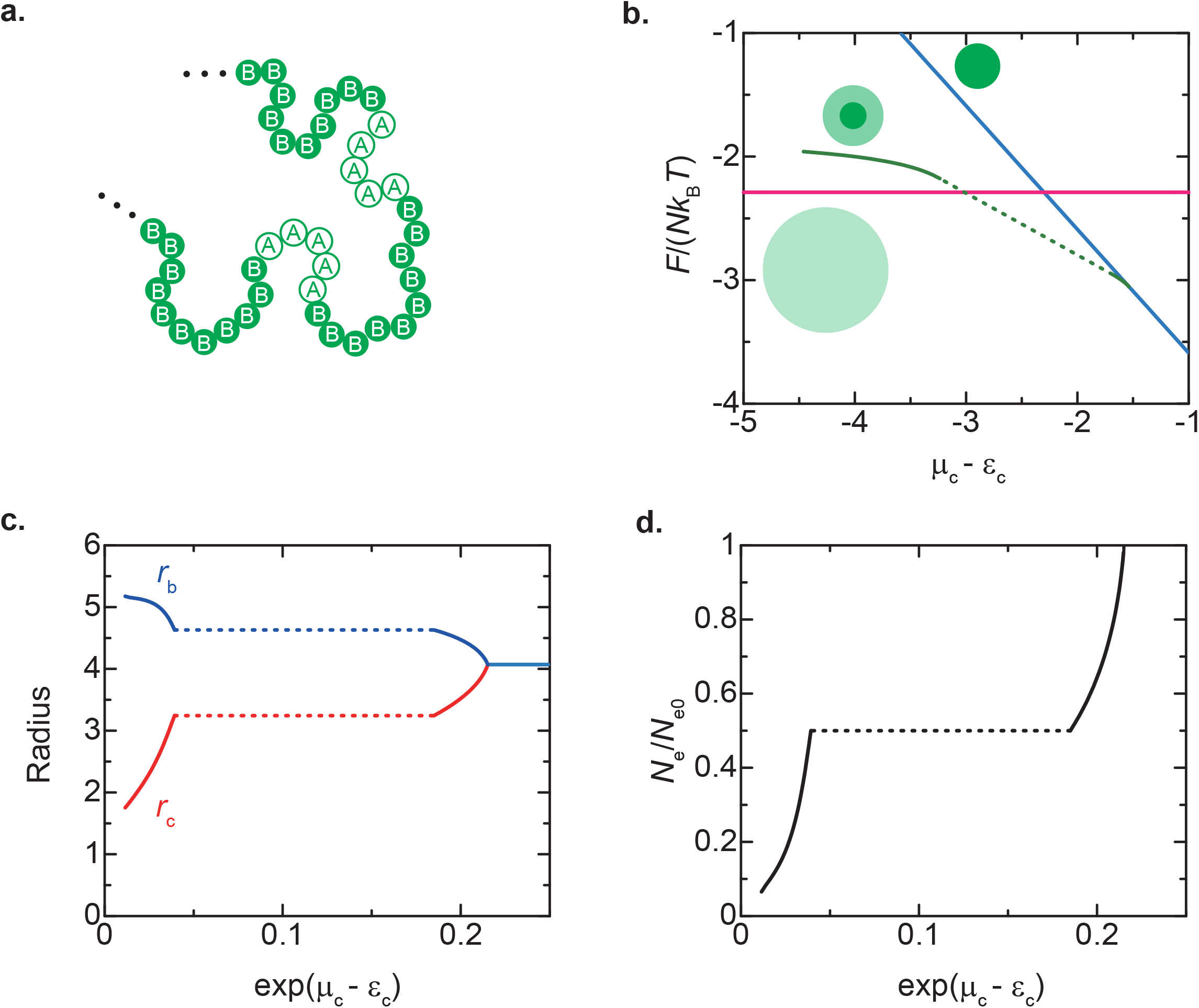
Multi-block copolymer model. **a**. DNAs are modeled as multi-block AB copolymers, which are composed of A and B blocks. The fraction of DNA units in A blocks is *N*_A_/*N* and the fraction of DNA units in B blocks is 1 − *N*_A_/*N*. The free energy *ϵ*_hA_ to assemble nucleosomes at A blocks is larger than the free energy *ϵ*_hB_ to assemble nucleosomes at B blocks. The values of the other parameters of A and B blocks are equal and are shown in Table 1. **b, c**, and **d** are numerical calculations on multi-block DNAs with *N*_A_/*N* = 0.5, *ϵ*_hA_= 1.0, *ϵ*_hB_ = −2.0. **b**. The free energy of the entangled multi-block DNAs is shown as a function of the chemical potential *µ*_c_ of mutant condensin for the bean state (green), the uni-condensed state (cyan), and the uni-swollen state (magenta). **c**. The radii of the bean (red) and the core (blue), *r*_b_ and *r*_c_, are shown as a function of the exponential, exp(*µ*_c_ − *ϵ*_c_), which is proportional to the concentration of mutant condensins. **d**. The fraction *N*_e_/*N*_e0_ of DNA units in the core is shown as a function of the exponential exp(*µ*_c_ − *ϵ*_c_). The fraction *N*_e_/*N*_e0_ does not change at a window of mutant condensin concentrations, shown by the dotted lines.

The multi-block copolymer model predicts that the bean structure can be most stable in a window of mutant condensin concentration if the difference *ϵ*_hA_ − *ϵ*_hB_ of the free energy of nucleosome assembly is large enough, see Fig. 8**b**. The feature of the bean structure composed of multi-block DNAs is that in a finite window of mutant condensin concentration, the radii, *r*_b_ and *r*_c_, and the distribution of DNA units between the core and the halo do not change with changing the mutant condensin concentration, see the dotted lines in Fig. 8**c** and **d**. In this window of mutant condensin concentration, all the B blocks are localized at the halo and all the A blocks are localized at the core, see Fig. 8**d**.

## DISCUSSION

We have constructed a theoretical model to provide insight into the formation of the bean structure. Our model is constructed based on the assumption that DNA subchains become stifer when they form nucleosomes or when part of the DNA is reeled out into the halo. This stifening mechanism results from the slip-link nature of polymer entanglements and the entropic nature of DNA elasticity in the chromosome length scale. To date, only a few theoretical studies have considered the conversion of DNA elasticity by the assembly and disassembly of nucleosomes in the context of DNA brush (Yamamoto and Schiessel, 2016, Yamamoto and Schiessel, 2017**a** and **b**, Leidescher et al., 2020). We also assumed that nucleosomes and mutant condensins are mutually exclusive on DNA. Thus, rich crosstalk between nucleosomes and mutant condensins controls the elasticity of entangled DNAs. Such crosstalk has not been considered in previous polymer simulations of mitotic chromosome assembly (Naumova et al., 2013, Goloborodko et al., 2016a and 2016b).

Our theory predicts that the DNA loops in the halo are stabilized by the preferential enrichment of nucleosomes in this region. The formation of the halo increases the elastic free energy of entangled DNA in the core, the elastic free energy of forming DNA loops in the halo, and the free energy of the DNA-DNA excluded volume interaction in the halo. Our theory also predicts that the elastic free energy of entangled DNA in the core is greatly reduced by the accumulation of mutant condensins, which in turn suppress nucleosome assembly in that region. Despite its simplicity, our theory captures the essential features of the bean structure observed experimentally (Kinoshita et al., 2022), including the differential localization of nucleosomes and mutant condensins between the core and the halo, and the dependence of the bean and core radii on the mutant condensin concentration, see Figs. 5 and 6. Some more detailed features, such as the high concentration of mutant condensins at the interface between the halo and the core and the asymmetry of the increase and decrease of the core and bean radii, remain to be explained. Future studies should extend our current theory to explain such features.

Our current theory allows us to propose two models for bean assembly – the uniform polymer model and the multi-block copolymer model. On the one hand, if the DNAs are assumed to be uniform polymers in the chromosome length scale, the bean structure does not represent the global free energy minimum because of the elastic and interaction free energy contributions in the halo. In this case, the bean structure can only be observed if the entangled DNAs are compacted by mutant condensins prior to nucleosome assembly. This is indeed the case in the experiments using mouse sperm nuclei as a substrate in *Xenopus* egg extracts (Shintomi et al., 2017). This model can be tested by controlling the timing of the addition of mutant condensins. On the other hand, if the DNAs are assumed to be multi-block copolymers, where nucleosomes are preferentially assembled on one type of blocks, the bean structure may represent the global free energy minimum. In this case, the tendency of DNA regions that localize in the core and the halo depends on the underlying DNA sequences. Then, one might wonder how DNAs act as multi-block copolymers in the chromosome scale. In fact, in polymer physics, DNAs are usually considered as uniform polymers in such length scale based on the assumption that the DNA sequence is statistical (Marko and Siggia, 1995). However, there are genomic features of the chromosome length scale, such as the repeat sequence at the centromeres and telomeres and the competition between histone and transcription factor binding at the euchromatic regions (Joseph et al., 2017). The insights from our theory may be useful in designing future experiments to determine whether the bean is the most stable structure or is kinetically trapped.

Our theory is based on the assumption that the condensin-condensin interaction is multivalent, similar to the previous models (Sakai et al., 2018, Forte et al., 2022). The postulated condensin-condensin interaction is under the control of the SMC ATPase cycle (Kinoshita et al., 2022). However, such non-equilibrium switching of the condensin-condensin interaction only decreases the magnitude of the interaction if the interaction is multivalent, see eq. (18). Recently, it has been proposed that bivalent DNA-cohesin-DNA bridging interaction drives the condensation of DNA-cohesin complexes (Ryu et al., 2021). If the condensin-condensin interaction is assumed to be bivalent, then the non-equilibrium switching will have a more pronounced effect on chromosome structure. Clearly, more experimental data are needed to clarify this important issue and to further refine our current theory.

To extend our theory to the mechanism of mitotic chromosome assembly, it is necessary to consider the loop extrusion activity of condensin complexes and the strand-passing activity of topo II. If condensins localized at the core start to extrude loops, they will generate tension of the DNAs entangled in the core so that the DNAs in the halo are reeled into the core (Yamamoto and Schiessel, 2022). The magnitude of the interaction between wild-type condensin complexes may be smaller than the magnitude of the interaction between mutant condensin complexes because CAP-G subunits counteract with CAP-D2 subunits (Kinoshita et al., 2015). Consistent with this prediction, the wild-type condensin complexes form a so-called ‘entangled mass’, in which the halo and core are not structurally differentiated, under the same condition (Kinoshita et al, 2022). Topo II disentangles and individualizes chromosomes through its strand-passing activity (Pommier et al., 2016). The disentanglement of polymers results from the thermal motion of polymers slipping out of the effective crosslinks from the polymer ends – the reptation dynamics (Doi and Edwards, 1986, Likhtman and McLeish, 2002). The strand-passing activity of topo II can be viewed as transiently creating ‘ends’ in the middle of DNAs and thus can be treated by taking into account the reptation dynamics in an extension of our theory. The individualization of chromosomes changes the geometry of the system (from a concentrated or semi-dilute regime to a dilute regime) and a separate treatment is required.

In summary, our current theory proposes a new concept of the elasticity control of entangled DNAs by the crosstalk between nucleosomes and condensin. This theory predicts that the bean structure is kinetically trapped or that DNAs should be considered as multi-block copolymers. A better understanding of the biophysical properties of nucleosomal DNAs and condensins will be important to elucidate the mechanism of mitotic chromosome assembly. Such biophysical properties may be best studied by using a simple system, such as the bean structure discussed in this study.

## Data availability

The Mathematica file “beanmodelVer69(haloextension).nb” used for the numerical calculations is available in figshare with the identifier (https://doi.org/10.6084/m9.figshare.21517056).

