## Supplemental File for "Elasticity control of entangled chromosomes: crosstalk between condensin complexes and nucleosomes"

### S1 Minimization of the free energy

#### S1.1 Free energy

We here derive the conditions of the DNA volume fraction $\phi$, the occupancies, $\alpha_{\rm c}$ and $\alpha_{\rm h}$, of DNA by mutant condensins and nucleosomes, and the fraction $N_{\rm c}/N$ of DNA units in the core. The free energy of the bean structure is discussed in the main article. We here summarize it to be self-contained. The free energy of the bean structure is the sum of the free energy $F_{\rm c}$ of the core and the free energy $F_{\rm h}$ of the halo,

$$F = F_{\rm c} + F_{\rm h}. \tag{S1}$$

The free energy of the core has the form

$$F_{\rm c} = V_{\rm c} f_{\rm c}(\alpha_{\rm c}, \alpha_{\rm h}, \phi), \tag{S2}$$

where $V_{\rm c}$ is the volume of the core and $f_{\rm c}$ is the free energy density of the core. The free energy density $f_{\rm c}$ in the core has the form

$$f_{\rm c}(\alpha_{\rm c}, \alpha_{\rm h}, \phi) = f_{\rm ent} + f_{\rm sol} + f_{\rm bnd}. \tag{S3}$$

$f_{\rm ent}$ is the density of the elastic free energy due to the entanglement. $f_{\rm sol}$ is the density of the solution free energy due to the mixing entropy and the interaction between mutant condensins. $f_{\rm bnd}$ is the density of the binding free energy due to the assembly of nucleosomes and the loading of mutant condensins.

The free energy of the halo has the form

$$F_{\rm h} \quad = \quad \int_{r_{\rm c}}^{r_{\rm b}} dr 4\pi r^2 f_{\rm h}(\alpha_{\rm c}, \alpha_{\rm h}, \phi, r), \tag{S4}$$

where $\int_{r_\mathrm{c}}^{r_\mathrm{b}} dr 4\pi r^2$ is the volume integral over the halo. $r$ is the distance from the center of the bean structure. $r_\mathrm{c}$ is the radius of the core and $r_\mathrm{b}$ is the radius of the bean structure. $f_\mathrm{h}(\alpha_\mathrm{c}, \alpha_\mathrm{h}, \phi, r)$ is the free energy density of the halo and is a function of $\alpha_\mathrm{c}$, $\alpha_\mathrm{h}$, $\phi$, and $r$. The free energy density $f_\mathrm{h}(\alpha_\mathrm{c}, \alpha_\mathrm{h}, \phi, r)$ in the halo has the form

$$f_\mathrm{h}(\alpha_\mathrm{c}, \alpha_\mathrm{h}, \phi, r) = f_\mathrm{lop} + f_\mathrm{sol} + f_\mathrm{bnd}. \tag{S5}$$

$f_\mathrm{lop}$ is the elastic free energy due to the DNA looping at the halo. $f_\mathrm{sol}$ is the solution free energy and $f_\mathrm{bnd}$ is the binding free energy. $f_\mathrm{sol}$ and $f_\mathrm{bnd}$ are the same as the second and third terms in the round bracket of eq. (S2).

The volume $V_\mathrm{c}$ and the radius $r_\mathrm{c}$ of the core a relationship with the DNA volume fraction $\phi$ in the core

$$V_\mathrm{c} = \frac{4\pi}{3} r_\mathrm{c}^3 = \frac{b_0^3 N_\mathrm{c}}{\phi}, \tag{S6}$$

where $N_\mathrm{c}$ is the number of DNA units in the core. The radius $r_\mathrm{b}$ of the bean is determined by the conservation of the number of DNA units,

$$\int_{r_\mathrm{c}}^{r_\mathrm{b}} dr 4\pi r^2 \phi = b_0^3 (N - N_\mathrm{c}), \tag{S7}$$

where $N$ is the total number of DNA units in the system.

The elastic free energy has the form

$$f_\mathrm{ent} = \frac{3}{2} \frac{G_0}{\lambda_\mathrm{s}} \frac{N_\mathrm{e0}}{N_\mathrm{e}} \frac{1}{\alpha_\mathrm{h} + \rho(1 - \alpha_\mathrm{h})}. \tag{S8}$$

The derivation of eq. (S8) is shown in the main article and we do not repeat it here. $N_\mathrm{e0}$ and $N_\mathrm{e}$ are the number of DNA units between the effective crosslinks due to the entanglement in the relaxed (reference) state and the equilibrium (actual) state. $G_0$ is the shear modulus of the entangled chromosomes in the relaxed (reference) state, see eq. (13) in the main article. $\lambda_s$ is the swelling ratio. $\rho$ is the ratio between the size of bare DNA to the size of nucleosomal DNA, see eq. (14) in the main article. The number of effective crosslinks due to the entanglement is constant. Because the difference between $N_{e0}$ and $N_e$ is due to the assembly of DNA loops in the halo, the ratio $N_e/N_{e0}$ has a relationship

$$\frac{N_e}{N_{e0}} = \frac{N_c}{N}. \tag{S9}$$

The swelling ratio has a relationship with the DNA volume fraction,

$$\phi = \frac{\phi_0}{\lambda_s}\frac{N_e}{N_{e0}}. \tag{S10}$$

By using eq. (S10), eq. (S8) is represented as a function of the DNA volume fraction $\phi$ and the occupancies, $\alpha_c$ and $\alpha_h$, of DNA by mutant condensins and nucleosomes.

The solution free energy has the form

$$f_{sol} = \frac{k_B T}{b_0^3}\left[(1-\phi-\alpha_h\phi-\alpha_c\phi)\log(1-\phi-\alpha_h\phi-\alpha_c\phi)-\chi\alpha_c^2\phi^2\right]. \tag{S11}$$

$k_B$ is the Boltzmann constant and $T$ is the absolute temperature. $b_0$ is the Kuhn length of bare DNA. $\chi$ is the interaction parameter.

The binding free energy has the form

$$\begin{aligned} f_{bnd} = \frac{k_B T}{b_0^3}&\left[\alpha_c\log\alpha_c+\alpha_h\log\alpha_h+(1-\alpha_c-\alpha_h)\log(1-\alpha_c-\alpha_h)\right. \\ &\left.+(\epsilon_c-\mu_c)\alpha_c+(\epsilon_h-\mu_h)\alpha_h\right], \end{aligned} \tag{S12}$$

where $\epsilon_c$ is the energy gain due to the loading of mutant condensins and $\mu_c$ is the chemical potential of mutant condensins. $\epsilon_h$ is the energy gain due to the assembly of nucleosomes and $\mu_h$ is the chemical potential of histone proteins.

The elastic free energy $f_{\text{lop}}$ due to the DNA looping has the form

$$f_{\text{lop}} = \frac{3}{2}\left(\frac{r_{\text{m}}}{r}\right)^4 \frac{1}{b_0^3}\frac{1}{\phi}\frac{\rho}{\alpha_{\text{h}} + \rho(1-\alpha_{\text{h}})}, \tag{S13}$$

where $r_{\text{m}}$ is the length scale defined by

$$r_{\text{m}}^2 = \frac{n b_0^2}{4\pi}. \tag{S14}$$

#### S1.2 DNA volume fraction and occupancies in the halo

The free energy in the halo is a functional of the DNA volume fraction $\phi(r)$ and the occupancies, $\alpha_{\text{c}}(r)$ and $\alpha_{\text{h}}(r)$, of DNA by mutant condensins and nucleosomes at the halo. To ensure the conservation of the number of DNA units in the halo, eq. (S7), we use the Lagrange multiplier

$$\tilde{F} = F - \mu_{\text{d}}\left[\int_{r_{\text{c}}}^{r_{\text{b}}} dr 4\pi r^2 \phi - b_0^3(N - N_{\text{c}})\right] \tag{S15}$$

to the free energy, eq. (S1). We note that the second term of eq. (S15) is necessary because the number of DNA units in the halo depends on $N_{\text{c}}$, which is determined by the free energy minimization. $\mu_{\text{d}}$ is the chemical potential of DNA units.

The variation of the free energy with respect to $\phi(r)$, $\alpha_{\text{c}}(r)$, and $\alpha_{\text{h}}(r)$ has the form

$$\begin{aligned}\delta F &= \int_{r_{\text{c}}}^{r_{\text{b}}} 4\pi r^2 dr \left[\left(\frac{\partial}{\partial \phi} f_{\text{h}}(\alpha_{\text{c}}, \alpha_{\text{h}}, \phi, r) - \mu_{\text{d}}\right)\delta\phi + \left(\frac{\partial}{\partial \alpha_{\text{c}}} f_{\text{h}}(\alpha_{\text{c}}, \alpha_{\text{h}}, \phi, r)\right)\delta\alpha_{\text{c}}\right. \\ &\quad \left. + \left(\frac{\partial}{\partial \alpha_{\text{h}}} f_{\text{h}}(\alpha_{\text{c}}, \alpha_{\text{h}}, \phi, r)\right)\delta\alpha_{\text{h}}\right], \end{aligned} \tag{S16}$$

The profiles of $\phi$, $\alpha_{\text{c}}$, and $\alpha_{\text{h}}$ at the free energy minimum satisfy $\delta F = 0$ for any variations

$\delta\phi$, $\delta\alpha_{\mathrm{c}}$, and $\delta\alpha_{\mathrm{h}}$. This leads to the conditions of the free energy minimum

$$\frac{\partial}{\partial\phi} f_{\mathrm{h}}(\alpha_{\mathrm{c}}, \alpha_{\mathrm{h}}, \phi, r) = \mu_{\mathrm{d}} \tag{S17}$$

$$\frac{\partial}{\partial\alpha_{\mathrm{c}}} f_{\mathrm{h}}(\alpha_{\mathrm{c}}, \alpha_{\mathrm{h}}, \phi, r) = 0 \tag{S18}$$

$$\frac{\partial}{\partial\alpha_{\mathrm{h}}} f_{\mathrm{h}}(\alpha_{\mathrm{c}}, \alpha_{\mathrm{h}}, \phi, r) = 0 \tag{S19}$$

Eqs. (S32) - (S34) represent the fact that the chemical potentials of DNA units, mutant condensins, and histone proteins are uniform in the halo and lead to the profiles of $\phi(r)$, $\alpha_{\mathrm{c}}(r)$, and $\alpha_{\mathrm{h}}(r)$ in the halo.

Eq. (S17) can be integrated as

$$f_{\mathrm{h}}(\alpha_{\mathrm{c}}, \alpha_{\mathrm{h}}, \phi, r) - \mu_{\mathrm{d}}\phi = -\Pi_{\mathrm{h}}(r), \tag{S20}$$

where $\Pi_{\mathrm{h}}(r)$ is the integral constant, but is a function of $r$ because of the dependence of $f_{\mathrm{h}}$ on $r$. Eliminating $\mu_{\mathrm{d}}$ in eq. (S20) by using eq. (S17) leads to the form

$$\Pi_{\mathrm{h}}(r) \quad = \quad \phi^2 \frac{\partial}{\partial\phi}\left(\frac{f_{\mathrm{h}}(\alpha_{\mathrm{c}}, \alpha_{\mathrm{h}}, \phi, r)}{\phi}\right) \tag{S21}$$

and the integral constant $\Pi(r)$ is indeed the osmotic pressure. Because of the local equilibrium at the iterface between the halo and the exterior solution, the osmotic pressure is zero at $r = r_{\mathrm{b}}$

$$\Pi_{\mathrm{h}}(r_{\mathrm{b}}) = 0. \tag{S22}$$

Eq. (S22) is one of the boundary conditions of eqs. (S17) - (S19).

We note that eqs. (S17) - (S22) are also derived by directly enforcing the conservation of DNA units, eq. (S7), instead of using the Lagrange multiplier, the second term of eq. (S15). In such case, one has to take into account the fact that the exterior radius $r_{\mathrm{b}}$ of the bean changes by

$$\delta r_{\mathrm{b}} = \frac{1}{4\pi r_{\mathrm{b}}^2 \phi_{\mathrm{b}}} \int_{r_{\mathrm{c}}}^{r_{\mathrm{b}}} dr 4\pi r^2 \delta\phi. \tag{S23}$$

due to the variation of the DNA volume fraction $\delta\phi$. This indeed leads to the same form as eq. (S16) with

$$\mu_{\mathrm{d}} = \frac{f(\alpha_{\mathrm{cb}}, \alpha_{\mathrm{hb}}, \phi_{\mathrm{b}}, r_{\mathrm{b}})}{\phi_{\mathrm{b}}}, \tag{S24}$$

where the subscripts b represent the values of quantities at $r = r_{\mathrm{b}}$, i.e. $\alpha_{\mathrm{cb}} = \alpha_{\mathrm{c}}(r_{\mathrm{b}})$, $\alpha_{\mathrm{hb}} = \alpha_{\mathrm{h}}(r_{\mathrm{b}})$, $\phi_{\mathrm{b}} = \phi(r_{\mathrm{b}})$. Eq. (S24) is indeed equivalent to eq. (S22). However, we use the Lagrange multiplier, eq. (S15), that makes the following calculations simpler.

#### S1.3 DNA volume fraction and occupancies in the core

We denote the DNA volume fraction in the core as $\phi_{\mathrm{c}}$ to distinguish it from the DNA volume fraction in the halo. The DNA volume fraction $\phi_{\mathrm{c}}$, the occupancies of mutant condensins and nucleosomes, $\alpha_{\mathrm{c}}$ and $\alpha_{\mathrm{h}}$, in the core are derived by the minimization of the free energy, eq. (S15). At the minimum of the free energy, the partial derivatives of the free energy with respect to $\phi_{\mathrm{c}}$, $\alpha_{\mathrm{c}}$, and $\alpha_{\mathrm{h}}$ are zero. This leads to the conditions

$$\Pi_{\mathrm{c}} = \Pi_{\mathrm{h}}(r_{\mathrm{c}}) \tag{S25}$$

$$\frac{\partial}{\partial \alpha_{\mathrm{c}}} \left( \frac{f_{\mathrm{c}}(\alpha_{\mathrm{c}}, \alpha_{\mathrm{h}}, \phi)}{\phi} \right) = 0 \tag{S26}$$

$$\frac{\partial}{\partial \alpha_{\mathrm{h}}} \left( \frac{f_{\mathrm{c}}(\alpha_{\mathrm{c}}, \alpha_{\mathrm{h}}, \phi)}{\phi} \right) = 0, . \tag{S27}$$

where the osmotic pressure $\Pi_{\mathrm{c}}$ in the core is defined as

$$\Pi_{\mathrm{c}} = \phi_{\mathrm{c}}^2 \frac{\partial}{\partial \phi_{\mathrm{c}}} \left( \frac{f_{\mathrm{c}}(\alpha_{\mathrm{c}}, \alpha_{\mathrm{h}}, \phi_{\mathrm{c}})}{\phi_{\mathrm{c}}} \right). \tag{S28}$$

Eq. (S25) represents the continuity of the osmotic pressure at the interface between the halo and the core. Eqs. (S26) and (S27) represent the fact that the chemical potentials of mutant condensins and histone proteins are uniform in the core.

#### S1.4 Distribution of DNA

The fraction $n_\mathrm{e}$ $(= N_\mathrm{e}/N_\mathrm{e0})$ of DNA units between effective crosslinks is equal to the fraction of DNA units in the core, see eq. (S9). The fraction $n_\mathrm{e}$ also derived by the minimization of the free energy. The first derivative of the free energy with respect to the fraction $n_\mathrm{e}$ has the form

$$\mu_\mathrm{d} = \mu_\mathrm{e}, \tag{S29}$$

where $\mu_\mathrm{e}$ is the chemical potential of DNA units in the core

$$\mu_\mathrm{e} = \frac{\partial}{\partial n_\mathrm{e}}\left(\frac{F_\mathrm{c}}{N}\right) + \frac{\partial}{\partial n_\mathrm{e}}\left(\frac{4\pi r_\mathrm{c}^3}{3N}\right)\Pi_\mathrm{h}(r_\mathrm{c}). \tag{S30}$$

The second term of eq. (S30) is due to the fact that the volume of the core can change under the osmotic pressure $\Pi_\mathrm{h}(r_\mathrm{c})$ (one may find more explicit if one note that $N_\mathrm{c}$ $(= Nn_\mathrm{e})$, see eq. (S9)). By using eqs. (S2), (S6), and (S25), eq. (S30) is represented by only the quantities of the core

$$\mu_\mathrm{e} = \frac{\partial}{\partial n_\mathrm{e}}\left(n_\mathrm{e}\frac{f_\mathrm{c} b_0^3}{\phi_\mathrm{c}}\right) + \frac{\Pi_\mathrm{c} b_0^3}{\phi_\mathrm{c}}. \tag{S31}$$

#### S1.5 Explicit forms of the continuity equations

The explicit forms of the continuity equations of chemical potentials in the halo are derived by substituting eqs. (S5) into eqs. (S17) - (S19),

$$\alpha_\mathrm{c}\log\alpha_\mathrm{c} + \alpha_\mathrm{h}\log\alpha_\mathrm{h} + (1-\alpha_\mathrm{c}-\alpha_\mathrm{h})\log(1-\alpha_\mathrm{c}-\alpha_\mathrm{h}) + (\epsilon_\mathrm{c}-\mu_\mathrm{c})\alpha_\mathrm{c} + (\epsilon_\mathrm{h}-\mu_\mathrm{h})\alpha_\mathrm{h}$$

$$-(1+\alpha_\mathrm{c}+\alpha_\mathrm{h})(\log(1-\phi-\alpha_\mathrm{h}\phi-\alpha_\mathrm{c}\phi)+1)-2\chi\alpha_\mathrm{c}^2\phi$$
$$-\frac{3}{2}\left(\frac{r_\mathrm{m}}{r}\right)^4\frac{1}{\phi^2}\frac{\rho}{\alpha_\mathrm{h}+\rho(1-\alpha_\mathrm{h})}=\mu_\mathrm{d} \tag{S32}$$
$$\log\alpha_\mathrm{c}-\log(1-\alpha_\mathrm{c}-\alpha_\mathrm{h})+(\epsilon_\mathrm{c}-\mu_\mathrm{c})-\log(1-\phi-\alpha_\mathrm{c}\phi-\alpha_\mathrm{c}\phi)-1$$
$$-2\chi\alpha_\mathrm{c}\phi=0 \tag{S33}$$
$$\log\alpha_\mathrm{h}-\log(1-\alpha_\mathrm{c}-\alpha_\mathrm{h})+(\epsilon_\mathrm{h}-\mu_\mathrm{h})-\log(1-\phi-\alpha_\mathrm{c}\phi-\alpha_\mathrm{c}\phi)-1$$
$$+\frac{3}{2}\left(\frac{r_\mathrm{m}}{r}\right)^4\frac{1}{\phi^2}\frac{\rho(\rho-1)}{(\alpha_\mathrm{h}+\rho(1-\alpha_\mathrm{h}))^2}=0. \tag{S34}$$

The osmotic pressure $\Pi_\mathrm{h}$ is derived by substituting eq. (S5) into eq. (S21) as

$$\begin{aligned}\Pi_\mathrm{h} &= -\log\left(1-\phi-\alpha_\mathrm{c}\phi-\alpha_\mathrm{h}\phi\right)-(1+\alpha_\mathrm{c}+\alpha_\mathrm{h})\phi-\chi\alpha_\mathrm{c}^2\phi^2\\ &\quad -3\left(\frac{r_\mathrm{m}}{r}\right)^4\phi^{-1}\frac{\rho}{\alpha_\mathrm{h}+\rho(1-\alpha_\mathrm{h})}.\end{aligned} \tag{S35}$$

Similary, the continuity equations of chemical potentials are derived by substituting eq. (S3) into eqs. (S26) and (S27). This leads to eq. (S33) and

$$\log\alpha_\mathrm{h}-\log\left(1-\alpha_\mathrm{c}-\alpha_\mathrm{h}\right)+\epsilon_\mathrm{h}-\mu_\mathrm{h}-\log\left(1-\phi-\alpha_\mathrm{c}\phi-\alpha_\mathrm{h}\phi\right)-1$$
$$+\frac{3}{2}\frac{G_0b_0^3}{\phi_0^{1/3}k_\mathrm{B}T}\left(\frac{N_\mathrm{e0}}{N_\mathrm{e}}\right)^{4/3}\phi^{-2/3}\frac{\rho-1}{(\alpha_\mathrm{h}+\rho(1-\alpha_\mathrm{h}))^2}=0. \tag{S36}$$

The osmotic pressure $\Pi_\mathrm{c}$ in the core is derived by substituting eq. (S3) into eq. (S28),

$$\begin{aligned}\Pi_\mathrm{c} &= -\log\left(1-\phi_\mathrm{c}-\alpha_\mathrm{h}\phi_\mathrm{c}-\alpha_\mathrm{c}\phi_\mathrm{c}\right)-(1+\alpha_\mathrm{h}+\alpha_\mathrm{c})\phi_\mathrm{c}-\chi\alpha_\mathrm{c}^2\phi_\mathrm{c}^2\\ &\quad -\frac{G_0b_0^3}{\phi_0^{1/3}k_\mathrm{B}T}\left(\frac{N_\mathrm{e0}}{N_\mathrm{e}}\right)^{4/3}\phi_\mathrm{c}^{1/3}\frac{1}{\alpha_\mathrm{h}+\rho(1-\alpha_\mathrm{h})}.\end{aligned} \tag{S37}$$

### S2 Approximate solutions for uniform polymer model

#### S2.1 Free energy for uni-condensed state and bean core

In the condensed phase, the occupancy of DNA by mutant condensins is approximately unity, $\alpha_{\rm c} \approx 1$ and $\alpha_{\rm h} \approx 0$, and the volume fraction of DNA is approximately 1/2, $\phi \approx 1/2$. The free energy of the core thus has an asymptotic form

$$\frac{F_{\rm c}}{Nk_{\rm B}T} \simeq \frac{N_{\rm e}}{N_{\rm e0}}\left[(\epsilon_{\rm c}-\mu_{\rm c})-\frac{1}{2}\chi+\frac{3}{2^{1/3}}\frac{G_0 b_0^3}{\rho\phi_0^{1/3}k_{\rm B}T}\left(\frac{N_{\rm e0}}{N_{\rm e}}\right)^{4/3}\right]. \tag{S38}$$

Eq. (S38) represents that the free energy contribution of the loading of mutant condensin (the first term), the free energy contribution of the condensin-condensin interaction (the second term), and the elastic free energy contribution due to the entanglement (the third term) are most dominant in the free energy of the condensed core. The free energy of the uni-condensed state is derived by setting $N_{\rm e}/N_{\rm e0} = 1$.

#### S2.2 Free energy for uni-swollen state

We here derive the free energy for the uni-swollen state when the shear modulus $G_0 b_0^3/(k_{\rm B}T\phi_0^{1/3})$ due to the entanglement is small. In such case, the occupancy of mutant condensins is approximately zero, $\alpha_{\rm c} \approx 0$, and the occupancy of nucleosomes is approximately unit, $\alpha_{\rm h} \approx 1$. We thus derive the occupancy of nucleosomes in the form

$$\alpha_{\rm h} = 1 - \delta\alpha_{\rm h}. \tag{S39}$$

We expand eqs. (S33), (S36), and (S37) with $\Pi = 0$ in a power series of $\alpha_{\rm c}$, $\delta\alpha_{\rm h}$, and $\phi$ and omit the higher order terms. This leads to the forms

$$\log\alpha_{\rm c} - \log(\delta\alpha_{\rm h} - \alpha_{\rm c}) + \epsilon_{\rm c} - \mu_{\rm c} - 1 = 0 \tag{S40}$$

$$-\log(\delta\alpha_{\rm h} - \alpha_{\rm c}) + \epsilon_{\rm h} - \mu_{\rm h} - 1$$

$$+\frac{3}{2}\frac{G_0 b_0^3}{k_\mathrm{B}T\phi_0^{1/3}}\phi^{-2/3}\frac{\rho-1}{(1+(\rho-1)\delta\alpha_\mathrm{h})^2} = 0 \tag{S41}$$

$$-\frac{G_0 b_0^3}{k_\mathrm{B}T\phi_0^{1/3}}\phi^{1/3}\frac{1}{1+(\rho-1)\delta\alpha_\mathrm{h}} + 2(1-2\chi_\mathrm{h})\phi^2 + \frac{8}{3}\phi^3 = 0. \tag{S42}$$

By using eq. (S42), the DNA volume fraction is derived in the form

$$\phi = \left(\frac{G_0 b_0^3}{2k_\mathrm{B}T\phi_0^{1/3}}\right)^{3/5}\frac{1}{(1+(\rho-1)\delta\alpha_\mathrm{h})^{3/5}} \tag{S43}$$

for $\chi_\mathrm{h} = 0$ (athermal solvent) and

$$\phi = \left(\frac{3}{8}\frac{G_0 b_0^3}{k_\mathrm{B}T\phi_0^{1/3}}\right)^{3/8}\frac{1}{(1+(\rho-1)\delta\alpha_\mathrm{h})^{3/8}} \tag{S44}$$

for $\chi_\mathrm{h} = 1/2$ (theta solvent). In the following, we treat the case of the athermal solvent. One can find that the last term of eq. (S41) is negligible in the asymptotic limit, $G_0 b_0^3/(k_\mathrm{B}T\phi_0^{1/3}) \ll 1$, by substituting eq. (S43) into eq. (S41). Eqs. (S40) and (S41) lead to the occupancies

$$\alpha_\mathrm{c} = \mathrm{e}^{-(\epsilon_\mathrm{c}-\mu_\mathrm{c})+(\epsilon_\mathrm{h}-\mu_\mathrm{h})} \tag{S45}$$

$$\delta\alpha_\mathrm{h} = \left(1+\mathrm{e}^{-(\epsilon_\mathrm{c}-\mu_\mathrm{c})+1}\right)\mathrm{e}^{(\epsilon_\mathrm{h}-\mu_\mathrm{h})-1}. \tag{S46}$$

The free energy of the uniform swollen state has an asymptotic form

$$\begin{aligned}\frac{F}{Nk_\mathrm{B}T} &\simeq \frac{3}{2}\frac{G_0 b_0^3}{k_\mathrm{B}T\phi_0^{1/3}}\phi^{-2/3}\frac{1}{1+(\rho-1)\delta\alpha_\mathrm{h}} + (\epsilon_\mathrm{h}-\mu_\mathrm{h}) - 2 + 2(1-2\chi_\mathrm{h})\phi \\ &\simeq (\epsilon_\mathrm{h}-\mu_\mathrm{h}) - 2 + 5\left(\frac{G_0 b_0^3}{2k_\mathrm{B}T\phi_0^{1/3}}\right)^{3/5}\frac{1}{(1+(\rho-1)\delta\alpha_\mathrm{h})^{3/5}},\end{aligned} \tag{S47}$$

where we used eqs. (S43) - (S46) to derive the last form of eq. (S47).

#### S2.3 Chemical potential at swollen-condensed transition

The chemical potential $\mu_{\rm c}^{\rm th}$ of mutant condensins at the transition between uni-swollen and uni-condensed is derived by the condition that the free energies of the uni-swollen and uni-condensed states are equal. This condition leads to the form

$$\mu_{\rm c}^{\rm th} \approx \left(\epsilon_{\rm c} - \frac{1}{2}\chi - (\epsilon_{\rm h} - \mu_{\rm h}) + 2\right) + \delta\mu_{\rm c}^{\rm ela}(\tilde{G}_0). \tag{S48}$$

with

$$\delta\mu_{\rm c}^{\rm ela}(x) = \frac{3}{2^{1/3}}x - \frac{5}{2}x^{3/5}\frac{1}{(\delta\alpha_{\rm h} + 1/\rho)^{3/5}}, \tag{S49}$$

where we denoted the rescaled shear modulus as

$$\tilde{G}_0 \equiv \frac{G_0 b_0^3}{\rho\phi_0^{1/3} k_{\rm B}T}. \tag{S50}$$

One can use the asymptotic form

$$\delta\alpha_{\rm h} = \left(1 + {\rm e}^{-\chi/2-(\epsilon_{\rm h}-\mu_{\rm h})+3}\right) {\rm e}^{(\epsilon_{\rm h}-\mu_{\rm h})-1}. \tag{S51}$$

for $\tilde{G}_0 \ll 1$.

#### S2.4 Chemical potentials of DNA units at bean-condensed transition

The thickness of the halo is zero, $r_{\rm b} = r_{\rm c}$, and the fraction of DNA units in the core is unity, $n_{\rm e} = 1$, at the maximum chemical potential $\mu_{\rm c}^{\rm max}$ of mutant condensins, at which the bean state is stable. The osmotic pressure is thus zero both at the halo and the core. By using eqs. (S31) and (S38) with $\Pi_{\rm c} = 0$, the chemical potential of DNA units at this condition is derived as

$$\mu_{\rm e} \simeq \epsilon_{\rm c} - \mu_{\rm c} - \frac{1}{2}\chi - \frac{1}{2^{1/3}}\frac{G_0 b_0^3}{\rho\phi_0^{1/3}k_{\rm B}T}. \tag{S52}$$

In the halo, the occupancy of nucleosomes is relatively large, $\alpha_{\rm h} \approx 1$, while the occupancy of DNAs by condensin is relatively small, $\alpha_{\rm c} \approx 0$. The DNAs in the halo are swollen due to the DNA-DNA excluded volume interaction and thus the DNA volume fraction is small, $\phi < 1/2$. We derive the occupancy of nucleosomes in the halo in the form

$$\alpha_{\rm h} = 1 - \delta\alpha_{\rm h} \tag{S53}$$

with $\delta\alpha_{\rm h} \ll 1$. The chemical potentials of mutant condensins and histones as well as the osmotic pressure in the halo have asymptotic forms

$$\log\alpha_{\rm c} - \log(\delta\alpha_{\rm h} - \alpha_{\rm c}) + (\epsilon_{\rm c} - \mu_{\rm c}) - 1 = 0 \tag{S54}$$

$$-\log(\delta\alpha_{\rm h} - \alpha_{\rm c}) + (\epsilon_{\rm h} - \mu_{\rm h}) - 1 + \frac{3}{2}\left(\frac{r_{\rm m}}{r_{\rm c}}\right)^4 \frac{\rho^2}{(1+(\rho-1)\delta\alpha_{\rm h})^2}\frac{1}{\phi^2} = 0 \tag{S55}$$

$$-3\left(\frac{r_{\rm m}}{r_{\rm c}}\right)^4 \frac{1}{\phi}\frac{\rho}{1+(\rho-1)\delta\alpha_{\rm h}} + 2\phi^2 = 0, \tag{S56}$$

where these relationships are derived by expanding eqs. (S33), (S34), and (S35) in a power series of $\delta\alpha_{\rm h}$, $\alpha_{\rm c}$, and $\phi$ and neglect the higher order terms with $\Pi_{\rm h} = 0$.

The occupancy $\alpha_{\rm c}$ of mutant condensins and the DNA volume fraction $\phi$ are derived in terms of $\delta\alpha_{\rm h}$

$$\alpha_{\rm c} = \frac{{\rm e}^{-(\epsilon_{\rm c}-\mu_{\rm c})+1}}{1+{\rm e}^{-(\epsilon_{\rm c}-\mu_{\rm c})+1}}\delta\alpha_{\rm h} \tag{S57}$$

$$\phi = \left(\frac{3}{2}\right)^{1/3}\left(\frac{r_{\rm m}}{r_{\rm c}}\right)^{4/3}\left(\frac{\rho}{1+(\rho-1)\delta\alpha_{\rm h}}\right)^{1/3}. \tag{S58}$$

by using eqs. (S54) and (S56). The radius $r_\mathrm{c}$ of the core is approximated as

$$\left(\frac{r_\mathrm{m}}{r_\mathrm{c}}\right)^4 = \left(\frac{\phi_\mathrm{m}}{\phi_\mathrm{c}} n_\mathrm{e}\right)^{-4/3} \simeq (2\phi_\mathrm{m})^{-4/3} \tag{S59}$$

by using $\phi_\mathrm{c} \approx 1$ and $n_\mathrm{e} = 1$. By using eq. (S55), $\delta\alpha_\mathrm{h}$ is derived as

$$\delta\alpha_\mathrm{h} = \frac{2^{5/4}}{3^{1/2}}(2\phi_\mathrm{m})^{-1/3}\left(\frac{1}{\mathrm{Plog}\left(\frac{4}{3}a^{4/3}\right)}\right)^{3/4}, \tag{S60}$$

where $\mathrm{Plog}(x)$ is the product-log function ($y = \mathrm{Plog}(x)$ is the solution of $x = y\mathrm{e}^y$) and $a$ is defined by

$$\log a = 1 - (\epsilon_\mathrm{h} - \mu_\mathrm{h}) + \log\left[\left(\frac{3}{2}\right)^{1/4}(2\phi_\mathrm{m})^{-1/3}\right] - \log\left(1 + \mathrm{e}^{-(\epsilon_\mathrm{c}-\mu_\mathrm{c})+1}\right). \tag{S61}$$

Eq. (S60) has an asymptotic form

$$\delta\alpha_\mathrm{h} = \left(\frac{3}{2}\right)^{1/4}(2\phi_\mathrm{m})^{-1/3}(1 - (\epsilon_\mathrm{h} - \mu_\mathrm{h}))^{-3/4} \tag{S62}$$

for negative large values of $\epsilon_\mathrm{h} - \mu_\mathrm{h}$.

The chemical potentials of DNA units has an asymptotic form

$$\mu_\mathrm{d} \simeq (\epsilon_\mathrm{h} - \mu_\mathrm{h}) - 2 + 3\phi + 4\phi^2, \tag{S63}$$

which is derived by expanding eq. (S32) with respect to $\delta\alpha_\mathrm{h}$, $\alpha_\mathrm{c}$, and $\phi$. For negative large values of $\epsilon_\mathrm{h} - \mu_\mathrm{h}$, the chemical potentials of DNA units have an asymptotic form

$$\mu_\mathrm{d} \simeq (\epsilon_\mathrm{h} - \mu_\mathrm{h}) - 2 + 3\left(\frac{3}{2}\right)^{1/4}(2\phi_\mathrm{m})^{-1/3}(1 - (\epsilon_\mathrm{h} - \mu_\mathrm{h}))^{1/4}. \tag{S64}$$

The equality of the chemical potentials of DNA units, $\mu_\mathrm{e} = \mu_\mathrm{d}$, leads to the chemical potential of mutant condensin at the bean-condensed transition in the form

$$\mu_{\mathrm{c}}^{\max} = \left(\epsilon_{\mathrm{c}} - (\epsilon_{\mathrm{h}} - \mu_{\mathrm{h}}) - \frac{1}{2}\chi + 2\right) - \frac{1}{2^{1/3}} \frac{G_0 b_0^3}{\rho \phi_0^{1/3} k_{\mathrm{B}} T} - 3\left(\frac{3}{2}\right)^{1/4} (2\phi_{\mathrm{m}})^{-1/3} (1 - (\epsilon_{\mathrm{h}} - \mu_{\mathrm{h}}))^{1/4}, \tag{S65}$$

see eq. (29) in the main article.

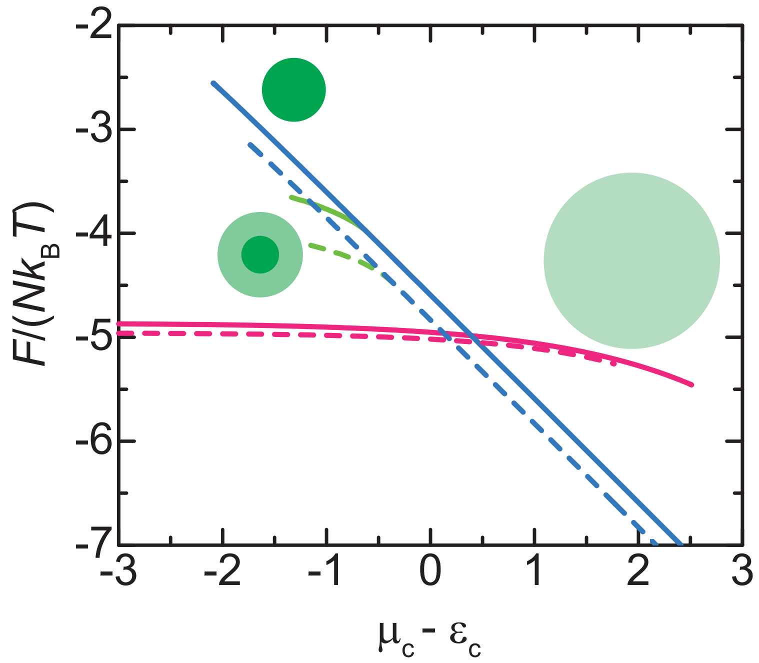


Figure S1: **Solvent dependence of free energy of entangled DNAs.** The free energy of entangled DNAs is shown as functions of the chemical potential $\mu_c - \epsilon_c$ of mutant condensins for the athermal solvent (solid) and $\theta$-solvent (broken) conditions. The values of the free energy of the bean, uni-condensed, and uni-swollen states are shown by the green, cyan, and magenta lines, respectively. The parameters used for the calculation were shown in Table 1.

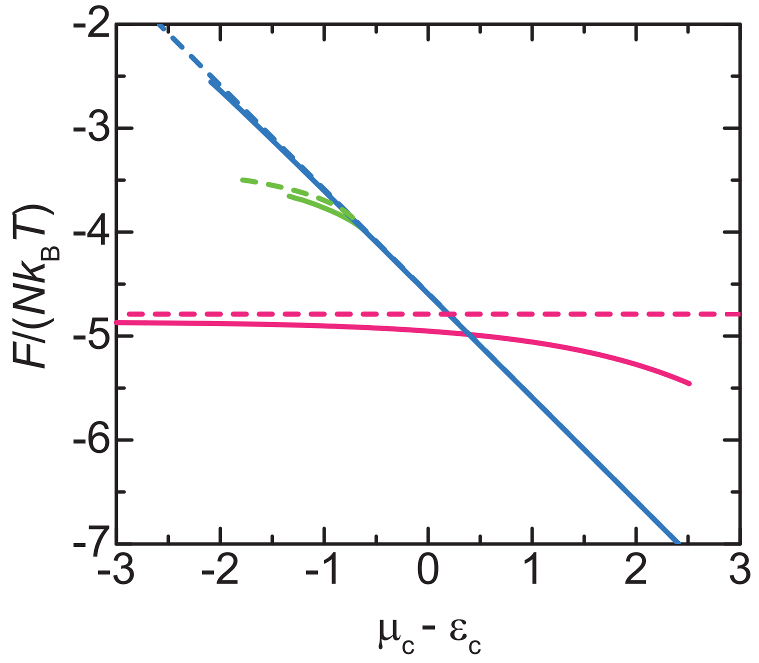


Figure S2: **Comparison between uniform polymer and multi-block copolymer models.** The free energy calculated by using the multi-block copolymer model (broken lines) for $N_A/N = 0$ and $\epsilon_{hB} = -3.0$, corresponding to the case of the uniform polymer model (solid), is shown as a function of the chemical potential $\mu_c$ of the mutant condensin concentration. The free energy was calculated for the bean (green), uni-condensed (cyan), and uni-swollen (magenta) states. The difference between the solid and broken lines results from the assumptions used for the multi-block copolymer model: $\alpha_c = 0$ in the halo and $\alpha_c = 1$ in the core in the bean state, $\alpha_c = 1$ in the uni-condensed state, and $\alpha_h = 1$ in the uni-swollen state.
